# IDENTIFICATION OF IMMUNE RESPONSE AND RNA NETWORK OF RHEUMATOID ARTHRITIS AND MOLECULAR DOCKING OF *CELASTRUS PANICULATUS* AS POTENTIAL THERAPEUTIC AGENT

**DOI:** 10.1101/2024.07.30.605947

**Authors:** Venkataramanan Swaminathan, Jeyvisna Arumainathan

## Abstract

**Background:** Rheumatoid Arthritis (RA) is an acute autoimmune disease leading to critical joint damage and bone destruction, weakening extra-articular organs over time. The pathogenesis of RA is complex and still undiscovered. This study aims to identify immune response, microRNA-hub genes network (miRNA), and drug candidates against RA via bioinformatics analysis.

**Methodology:** Three Gene Expression Omnibus (GEO) datasets were obtained from the NCBI database and classified into upregulated and downregulated differentially expressed genes (DEGs) using GEO2R tool. Gene enrichment analysis, protein-protein interaction network analysis, top 10 hub genes identification, miRNA-hub genes network analysis, and immune response identification were performed using various bioinformatic tools. Moreover, *Celastrus paniculatus* phytochemical compounds were retrieved and subjected to autodocking with upregulated and downregulated hub genes that are closely associated with RA. The drug-likeness and PreADMET analysis were performed.

**Results:** GSE30662, GSE766, GSE72100 datasets revealed 243 upregulated DEGs and 285 downregulated DEGs which exhibited RPS27A, UBB, UBC, UBA52, PSMD4, PSMD1, PSMD7, PSMB7, PSMD8, PSMA7 as top 10 upregulated hub genes and ACTB, TP53, AKT1, GAPDH, CTNNB1, EGFR, TNF, IL6, MYC, ANXA5 as top 10 downregulated hub genes. The miRNA network disclosed hsa-mir-23b-3p, as highly associated with upregulated hub genes whereas hsa-mir-34a-5p and hsa-mir-155-5p with downregulated hub genes. Additionally, immune responses of specific hub genes of RA were revealed while the docking analysis showed oleic acid and zeylasterone as novel drug candidates against RA.

**Conclusion:** Thus, hsa-mir-23b-3p, hsa-mir-34a-5p, and hsa-mir-155-5p can serve as therapeutic targets of RA while oleic acid and zeylasterone become potential drug candidates against RA.

## INTRODUCTION

Rheumatoid Arthritis (RA) is an acute inflammatory disease distinguished by inflammation of synovial, joint damage, and bone destruction. It affects any synovial joints of the body, starting with the small joints in the hands and feet (Black et al., 2023) Based on the Global Burden of Diseases, it is estimated that in 2020, 17.6 million people had RA and it is expected to elevate in 2050 whereby 31.7 million people will be affected by RA (Black et al., 2023). Women are 2-3 times more prone to RA than men due to several factors such as hormonal fluctuations, and the presence of a double X chromosome (He et al., 2022). In addition, people of any age can be affected by RA. However, adults aged 50-59 years old are more susceptible.

In an early stage of RA, the tissues around the joints are inflamed causing pain and stiffness. In the second stage, the inflammation begins to damage the cartilage of the joints (Luo et al., 2023). This will increase the stiffness and reduce motion. As RA progresses to the third stage, physical changes are also noticeable, the inflammation worsens the bones (Luo et al., 2023). This elevates pain, stiffness, and even less range of motion than in the second stage. Furthermore, in stage 4, the inflammation ends. However, there will be severe pain and swelling causing loss of mobility (Luo et al., 2023). Over time, it can also affect the heart, lungs, and nervous system causing fatal (Xing et al., 2022; Luo et al., 2023).

The signs and symptoms of RA include pain, stiffness, swelling, tenderness in joints, and fatigue (Luo et al., 2023). The inflammation in the joints and the destruction of bones and cartilage drive these signs and symptoms. The pathogenesis of RA is not entirely discovered. However, a complex interaction between genes and environmental factors is found to implicate in the pathogenesis of it (Firestein et al., 2017). The pathogenesis of RA begins years before it exhibits symptoms (Firestein et al., 2017). The genetic and environmental factors trigger synovial membrane inflammation and attract leukocytes into the affected tissue (Chahuan et al., 2023; Sharma & Goel, 2023). This causes the CD4+ helper T-cell activation and proliferation that produces inflammatory cytokines such as Tumour Necrosis Factor Alpha (TNF alpha), Interleukin-1 (IL1), Interleukin-17 (IL17), and, Interleukin-6 (IL6) (Admin & Admin, 2014; Chahuan et al., 2023). The activated T-cells initiate the differentiation of B-cells into plasma cells to synthesise autoantibodies such as Rheumatoid Factor (RF) and antigen-presenting cells antigen (ACPA) (Radu & Bungau, 2021). These autoantibodies do not recognize the self-tissues and begin to target them. Then, the osteoclasts and the Matrix metalloproteinase (MMP) induced by cytokines cause cartilage and bone degradation (Admin & Admin, 2014; Chauhan et al., 2023). Eventually, the joint gets damaged and recruits neutrophils, T-cells, and B-cells into the affected area leading to the synovial membrane hyperplasia and angiogenesis (Admin & Admin, 2014). Consequently, the joint tissue such as cartilage, ligaments, and subchondral bone will be enzymatically destroyed. The severe RA will affect the whole body.

Thus, this proves that RA has a close connection with the immune system because the immune system of an RA patient mistakenly attacks its body tissues, especially synovium, resulting in inflammation, pain, and joint damage (Li et al., 2022). Besides, immune cells like monocytes, macrophages, B cells, and CD4+ T cells play an essential role in RA’s progression and inflammatory responses (Feng, 2023; VASANTHI et al., 2007). The inflammatory autoimmune response triggers the synthesis of autoantibodies such as RF and ACPA, contributing to joint inflammation and systemic complications (Yap et al., 2018).

RA is caused by abnormalities in the immune system that are influenced by genetic and environmental factors such as family history, smoking, highly industrialised countries, exposure to environmental toxins, and stress (He et al., 2022). Some of the main genetic factors associated with RA are the presence of risk alleles such as Human Leukocyte Antigen (HLA), tyrosine phosphatase non-receptor type-22 (PTPN22) alleles, tumours necrosis factor receptor-associated factor 1, and interferon regulatory 5 (IL-5) (Radu & Bungau, 2021). Furthermore, the diagnosis of RA includes blood tests and imaging tests. The blood test allows us to identify the inflammation and antibodies that are the signs of RA such as C-reactive protein (CRP), erythrocyte sedimentation rate (ESR), and rheumatoid factor (RF) (Luo et al., 2023). On the other hand, the imaging tests include X-ray, ultrasound, and MRI scans used to view the affected joints.

The current treatment for RA is intended to reduce the pain and swelling of joints as RA is not curable. The treatment includes lifestyle changes, physical therapy, medication, and surgery (Luo et al., 2023). Medications such as Disease-Modifying-Antirheumatic-Drug (DMARD), non-steroidal anti-inflammatory drugs (NSAIDs), and corticosteroids help to alleviate pain and slow down the progression of RA (Luo et al., 2023). In a very severe stage of RA, surgeries are performed to repair or replace the destructed joint. Despite these treatments, a remarkable percentage of RA patients still suffer from the adverse effects of the medications, disability, and reduced quality of life (Peng et al., 2023)Thus, the identification of new effective therapeutic approaches has become an urgent focal issue.

Bioinformatics analysis is becoming a beneficial tool to determine the hub genes, miRNA, and immune response associated with RA and discover potential drug candidates for it. Hub genes and immune responses are essential in understanding the underlying pathogenesis of RA and discovering shared pathophysiology of other related diseases. Besides, miRNA is a small and non-coding Ribonucleic Acid (RNA) molecule, responsible for cell proliferation, differentiation, signal transduction, and gene regulation post-transcriptionally (O’Brien et al., 2018; Zhang et al., 2023). Dysregulation of miRNA such as abnormal expression is closely related to the progression of RA. Thus, miRNAs are considered biomarkers for the prognosis and diagnosis of RA (Zhang et al., 2023).

In addition, Structure-Based Drug Design (SBDD) has paved the way to speed up the discovery and development of drug processes by identifying potential compounds to bind at the active site of the targeted protein (Salum et al., 2008). This approach has been extensively used to predict the drug candidates against various diseases including Severe-Acute-Respiratory-Syndrome-Related-CoronaVirus (SARS-COV-2) main protease inhibition by *Andrographis paniculata* (Enmozhi et al., 2020) and glycogen phosphorylase activity reduction by *Tinospora cordifolia* (Herowati & Widodo, 2014). Hence, an SBDD such as molecular docking is a promising tool to predict the potent drug candidates against chronic inflammatory diseases such as RA.

In this study, *Celastrus paniculatus (C.paniculatus)* was selected to evaluate the tendency to act as a potential therapeutic agent against RA. *C.paniculatus* is recognized as the Black Oil Plant which is a woody climbing shrub (Sharma et al., 2021). It belongs to the Celastraceae family and is native to the Indian subcontinent and is also widely distributed in many other countries such as Maldives, Malaysia, Taiwan, China, Japan, and Australia which makes it easier to find (Sharma et al., 2021). It has been extensively used in Traditional Chinese Medicine (TCM) and Indian Traditional Medicine (ITM), including Ayurveda, Unani, Siddha, and Sowa Rigpa to treat various ailments such as cognitive dysfunction, insomnia, skin problems, and neurologic disorders (Sharma et al., 2021). It is called Jyotishmati and Malkagni in Ayurveda (Kothavade et al., 2015). In TCM, the Celastrus species is used as functional food such as nervine tonics and healthy drinks. A study has mentioned that the seed oil of *C.paniculatus* can treat inflammation, paralysis, leprosy, fevers, headache, and depression, proving their potential to treat symptoms of RA (Shen et al., 2019).

*C.paniculatus* is composed of several phytochemical compounds that have therapeutic properties (Kothavade et al., 2015; Shen et al., 2019). The major active phytochemicals of *C.paniculatus* include sesquiterpene alkaloids, triterpenoids, fatty acids, and polyalcohol esters (Kothavade et al., 2015; Shen et al., 2019). There has been a remarkable rise in the biological interest in phytochemical compounds of Celastrus species recently (Shen et al., 2019). Nevertheless, there is a lack of literature information about the potential of *C.paniculatus* compounds as drug candidates for RA. Hence, this research aims to identify the hub genes, immune response, and miRNA-hub genes network associated with RA using bioinformatics analysis and discover potential compounds of *C.paniculatus* against RA through molecular docking. It becomes fundamental insights into the molecular immune mechanism and unfolds potent novel approaches for the treatment and management of RA.

## METHODOLOGY

### Gene Expressed Omnibus (GEO) Datasets Selection

The Gene Expressed Omnibus (GEO) datasets of RA were obtained from the National Center for Biotechnology Information (NCBI) (*National Center for Biotechnology Information*, 1988). They were obtained under the GEO database and the results were refined to an expression profiling array and particularized to homo sapiens species.

### Differentially Expressed Genes (DEGs) Identification

The Differentially Expressed Genes (DEGs) were filtered using the GEO2R analysis tool. The DEGs were identified based on a p-value of less than 0.05 and log fold change of more than 1 as it is the most effective parameter (Luo et al., 2023; Swaminathan et al., 2023). The upregulated and downregulated DEGs obtained from this analysis were classified based on log fold change positive and negative values. The Venn diagram analysis was also performed using an online tool, Bioinformatics and Genomics to identify the intersections of DEGs in the three datasets.

### Functional Enrichment Analysis

The Database for Annotation, Visualisation, and Integrated Discovery (DAVID) (DAVID Functional Annotation Bioinformatics Microarray Analysis, n.d.) was used to obtain the biological functions, pathways, and ontology of DEGs. The organism was specified for homo sapiens and a P-value less than 0.05 was set. The results of the top 10 entities for biological process (BP), molecular function (MF) cellular component (CC), and KEGG pathways were selected for both upregulated and downregulated DEGs. Moreover, the visualizations of all the GO terms and KEGG pathways were conducted using SRplot (Tang et al., 2023). The -log(p-value) was used to visualize the result in the barplot.

### Protein-Protein Interaction Network Analysis

STRING online database (STRING, 2021) was used to create a Protein-Protein Interaction (PPI) network of the upregulated and downregulated genes. The cut-off value of more than 0.4 was set for a medium confidence interaction score to prevent protein network errors.

### Identification of Hub Genes

Cytoscape (Ono, n.d.) is a software to visualize networks by various plugins. Cytoscape v 3.9.1 was installed and cytohubba was plugged in using the App Manager. The top 10 hub genes were identified for upregulated and downregulated genes respectively by setting maximum interactions to 10 and confidence cutoff value to 0.4. The hub genes networks were obtained using the Maximal Clique Centrality (MCC) method, which has the highest performance for protein-to-protein networks (Luo et al., 2023).

### MiRNA-Hub Genes Network Analysis

miRnet2.0 (*MiRNet*, n.d.) was used to design and evaluate the miRNA-hub genes network for the top 10 upregulated and downregulated hub genes associated with RA. The organisms were specified for homo sapiens, selected the official gene symbol option, and uploaded the upregulated and downregulated list of hub genes separately in miRnet 2.0. miRNA nodes with 3 degrees and above were analyzed using a degree filter to get more significant results.

### Identification Of Immune Response

The InnateDB (Al, n.d.) is an open resource that acts as a knowledgebase to identify and visualize the innate immune response of humans, mice, and bovines. It is used to identify the immune response of specific hub genes that are highly associated with RA for upregulated and downregulated hub genes. The organisms were set as homo sapiens, the gene name was entered, and the return Genes option was selected.

### Retrieval of *Celastrus Paniculatus* Compounds

The phytochemical compounds present in *C.paniculatus* were retrieved from Indian Medicinal Plants, Phytochemistry and Therapeutics (IMPPAT) (IMPPAT, n.d.) A total of 20 phytochemical compounds of *C.paniculatus* were obtained from the database. Plant information such as taxonomy classification, common names, and systems of medicine were also gathered from this database.

### Selection and Preparation of Ligands

The three-dimensional structures of 20 compounds of *C. Paniculatus* were downloaded from PubChem (PubChem, 2004) in SDF format. It was converted to PDB format using Pymol software and then subjected to Autodock Vina v1.5.7. The compounds were set up in Autodock Vina v1.5.7 by adding gasteiger charges, merging non-polar hydrogens, and setting TORSDOF to obtain the most stable conformation (Al-Khayri et al., 2023). The setup ligands were then saved in PDBQT format.

### Selection and Preparation of Targeted Proteins

The PDB structures of the highly associated upregulated and downregulated proteins with RA were downloaded from the RCSB Protein Data Bank (Bank, n.d.) database. The preparation of targeted proteins was done using Autodock Vina v1.5.7 by removing water molecules, adding hydrogen bonds to polar sides, and adding default Kollman charges to the proteins. The protein was saved in PDBQT format.

### Autodocking

The autodocking was executed using Autodock Vina v1.5.7 to identify the potential drug candidates against the highly associated upregulated and downregulated proteins with RA. The 20 ligands were docked with upregulated and downregulated targeted proteins that are closely related to RA, and each docking result’s binding affinities were tabulated. The molecular interactions of ligands and target proteins with the best binding affinities were visualised using Pymol and Discovery Studio.

### Drug-likeness and Pre-Admet Analysis

SwissADME (SwissADME, n.d) and ADMETlab 2.0 (*ADMETLab 2.0*, n.d.) are web servers used to perform predictions of drug-likeness, pharmacokinetics properties, absorption, distribution, metabolism, excretion, and toxicity (ADMET) of the drug candidates to speed up drug discovery as it helps to eliminate undesirable drugs easily. The drug-likeness and PreADMET analysis for best binding affinity compounds were performed using the SwissADME and ADMETlab 2.0 web servers respectively by submitting the compounds in SMILE format.

## RESULTS

### Gene Expression Profiling of RA

This study evaluates the essential GEO datasets to comprehend gene activities and obtain information about the relationship between genes of various microarray datasets. Based on Table 1, GSE30662, GSE766, and GSE72100 were selected. They contain 8 samples, 6 samples, and 6 samples respectively. Each sample ID of the datasets has its unique GSE number and contains sample sizes less than 10. Moreover, in all three 3 datasets, various treatments were carried out such as Leukocytaphersis, TNF-alpha, and biological agents to investigate their efficacy in treating RA. The GEO2R analysis tool was used to identify the upregulated and downregulated genes using log 2 fold change more than 1.0 and p-value less than 0.05. In the GSE30662 dataset, 7435 DEGs were identified (3730 upregulated and 3705 downregulated) while GSE766 has 5432 DEGs (2378 upregulated and 3054 downregulated). Moreover, GSE727100 contains 22599 DEGs (10628 upregulated and 11971 downregulated).

**Table 1:** Information on the selected datasets associated with RA.

| GEO Accession | Total Sample | Platform | Experiment Type |
| --- | --- | --- | --- |
| GSE30662 | 8 | GPL5639 | Expression Profiling by Array |
| GSE766 | 6 | GPL8300 | Expression Profiling by Array |
| GSE72100 | 6 | GPL17077 | Expression Profiling by Array |

According to the Venn Diagram analysis, 243 DEGs had upregulated expressions while 285 DEGs had downregulated expressions shared between the three datasets (Figure 2).

**Figure 1:**
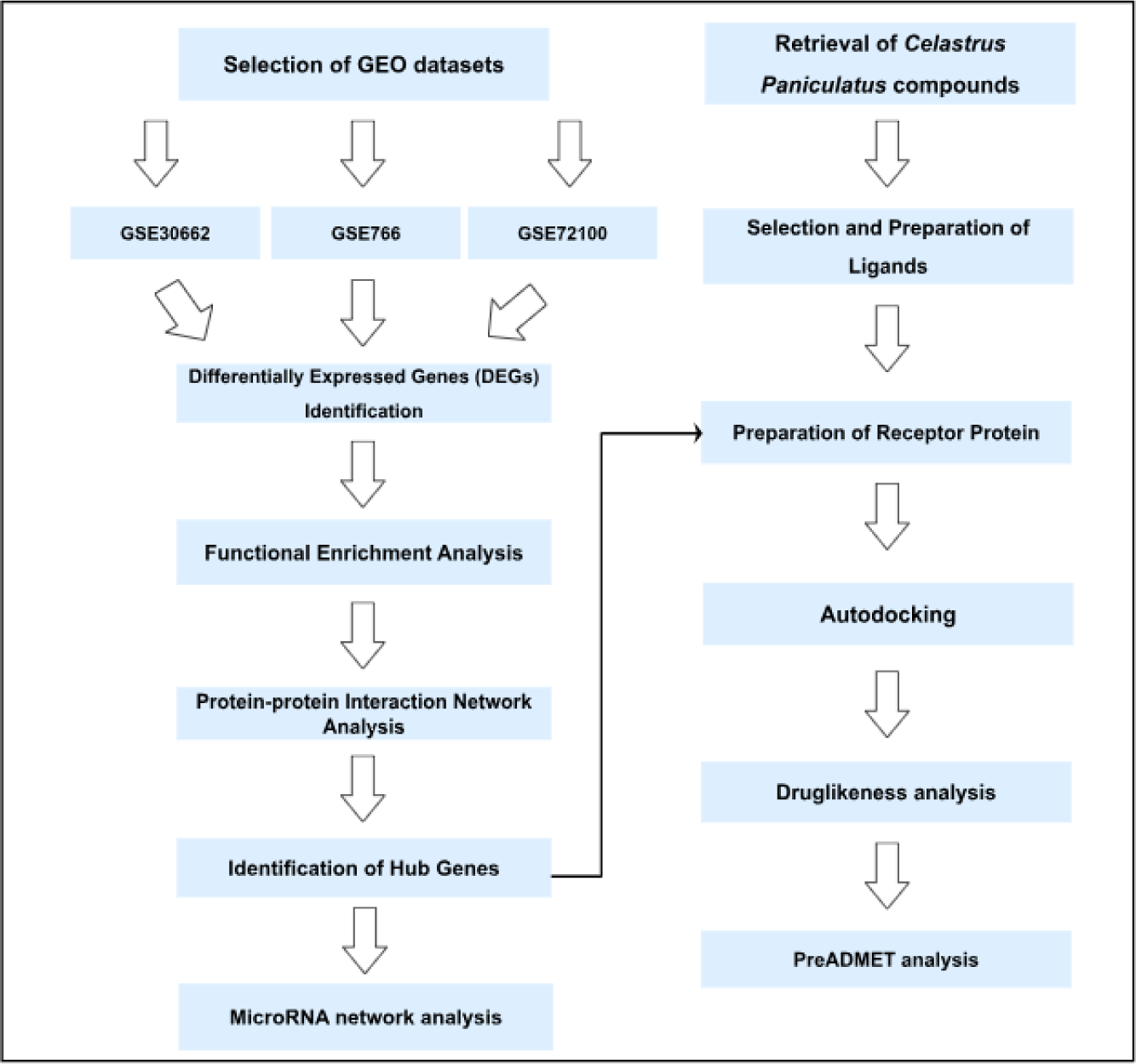
Flowchart of the research.

**Figure 2:**
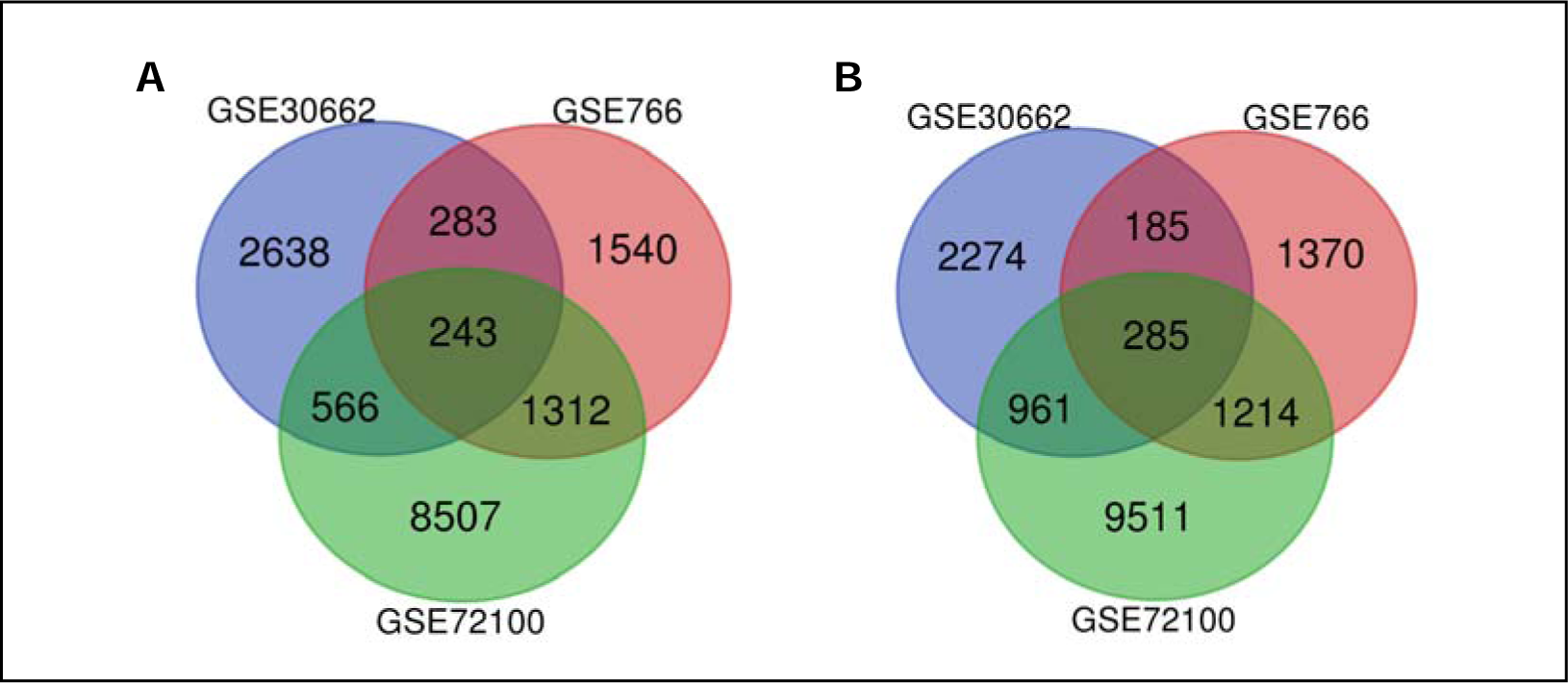
Identification of intersected DEGs among GSE30662, GSE766, and GSE72100 datasets. **(A)** DEGs with upregulated genes for all three datasets. **(B)** DEGs with downregulated genes for all three datasets.

### Functional Enrichment Analysis of Differentially Expressed Genes (DEGs)

The functional enrichment analysis was performed to explore the biological functions and enriched pathways of the DEGs. It was discovered that the DEGs were enriched in various processes and pathways such as biological processes (BP), molecular functions (MF), cellular components(CC), and Kyoto Encyclopedia of Genes and Genome (KEGG) pathways. The top 10 most enriched processes and pathways for upregulated and downregulated are listed in Table 2, Table 3, Table 4, and Table 5. Moreover, the visualization is depicted in Figure 3 and Figure 4. The functional enrichment analysis revealed that upregulated DEGs are mainly enriched in ubiquitin-dependent protein catabolic process, proteasome-mediated ubiquitin-dependent protein catabolic, protein stabilization, transcription from RNA polymerase II promoter, and mRNA splicing via spliceosome for the BP. In contrast, the upregulated DEGs are also notably associated with RNA, mRNA, protein, and DNA binding and transcription coactivator activity for MF. The nucleus, cytosol, nucleoplasm, proteasome complex, and cytoplasm are some of the enriched cellular components of upregulated DEGs.

**Figure 3:**
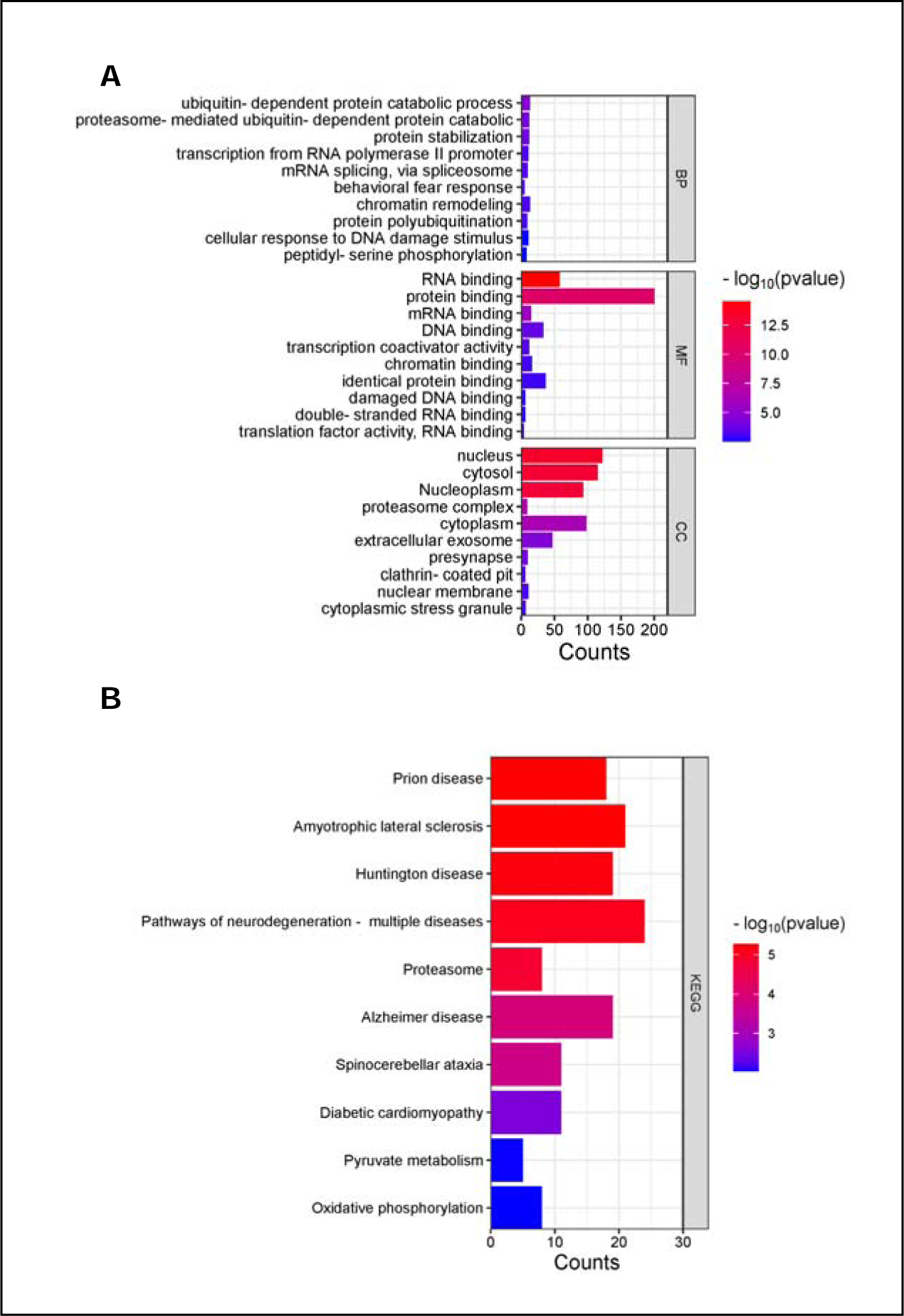
Visualization of gene enrichment analysis for upregulated hub genes. **(A)** GO terms and **(B)** KEGG analysis of the upregulated hub genes. Abbreviations: BP: Biological Process, MF: Molecular Function, CC: Cellular Components, KEGG: Kyoto Encyclopedia of Genes and Genome, DNA: Deoxyribonucleic Acid, RNA: Ribonucleic Acid

**Figure 4:**
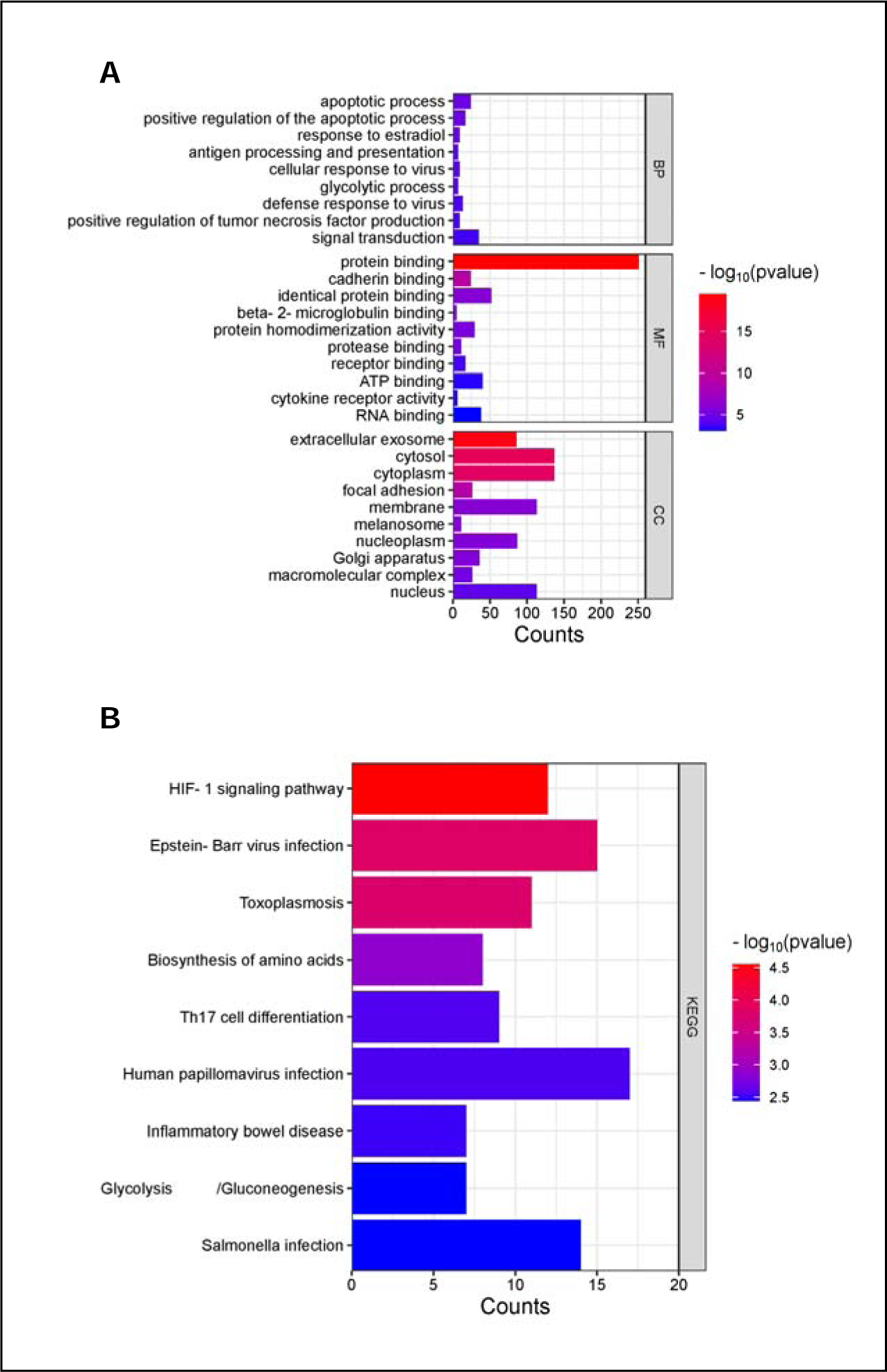
Visualization of gene enrichment analysis for downregulated hub genes. **(A)** GO terms and **(B)** KEGG analysis of the downregulated hub genes. Abbreviations: BP: Biological Process, MF: Molecular Function, CC: Cellular Components, KEGG: Kyoto Encyclopedia of Genes and Genome, RNA: Ribonucleic Acid, HIF-1: Hypoxia Inducible Factor-1, Th17: T-helper 17

**Table 2:**
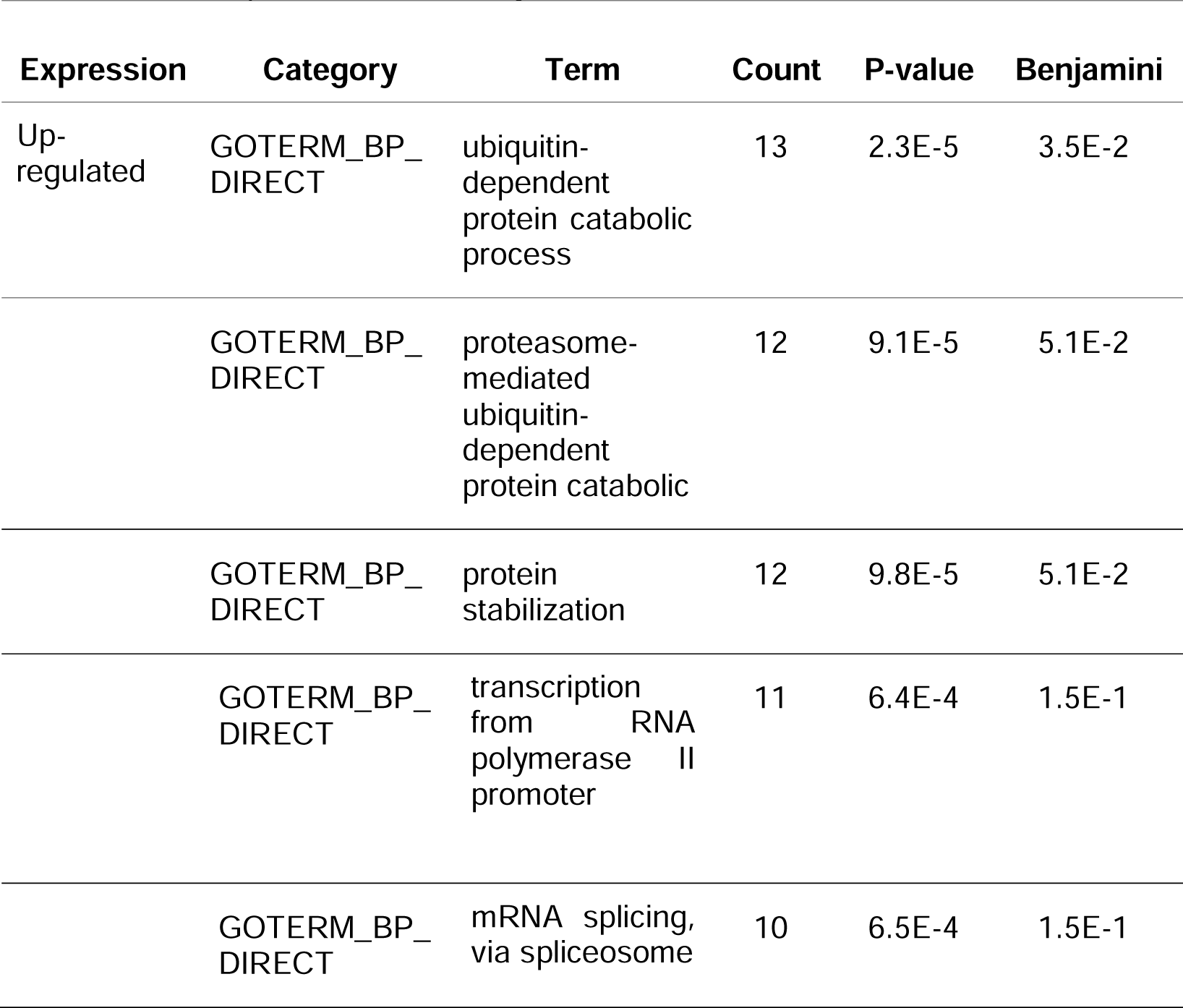
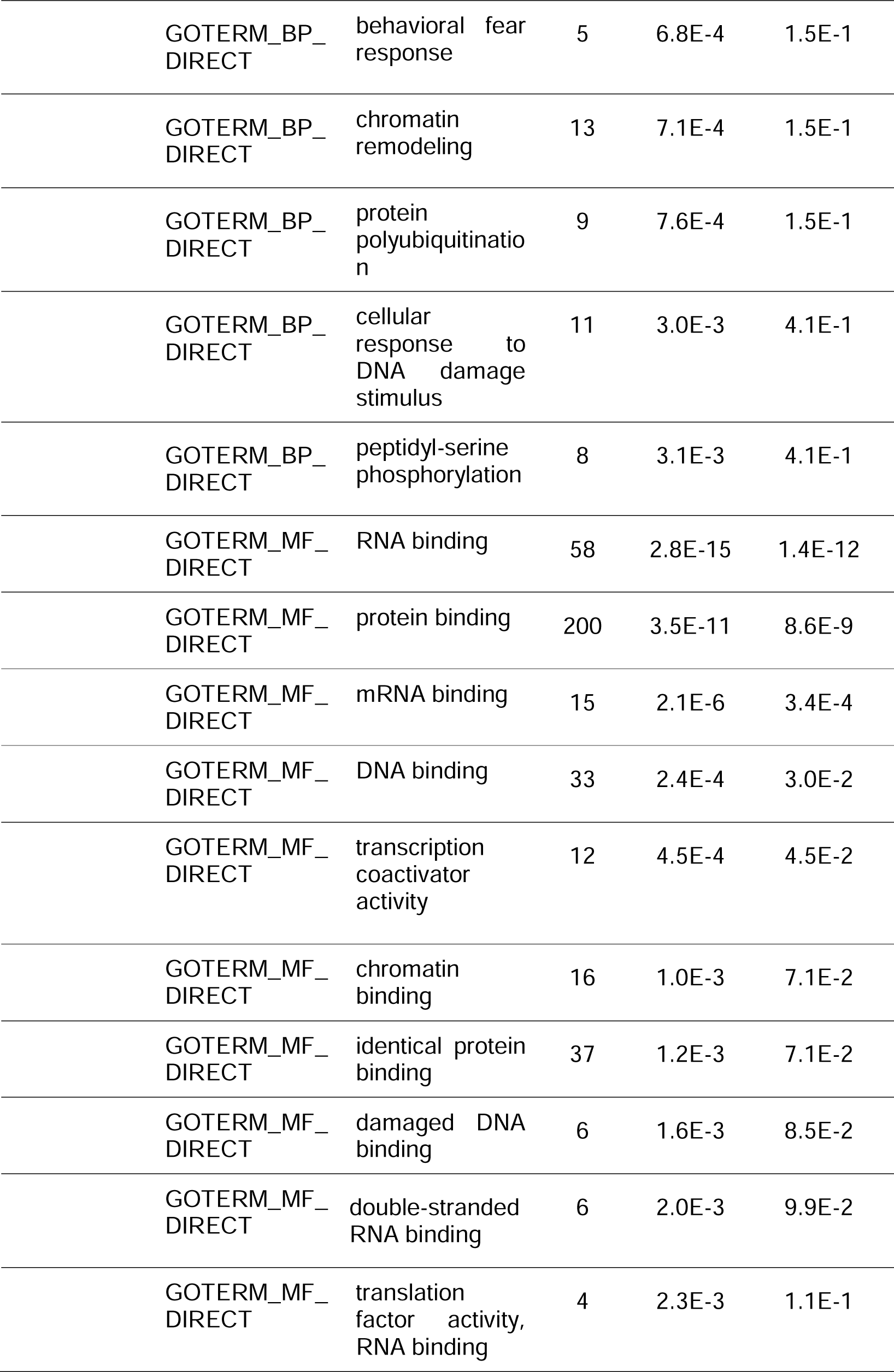
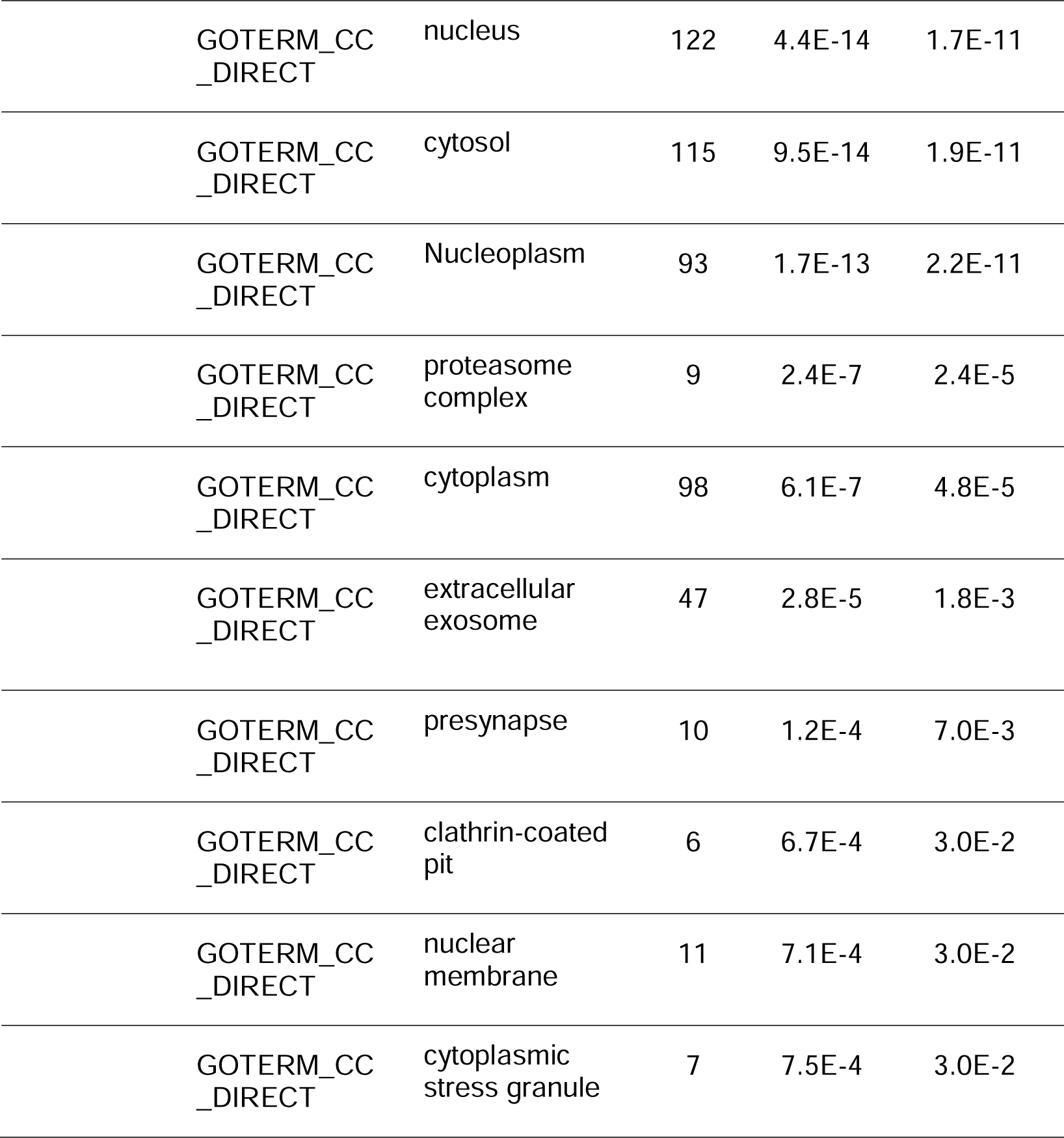
Top 10 Gene Ontology of upregulated DEGs of RA. Abbreviations: GOTERM: Gene Ontology Terms, BP: Biological Process, MF: Molecular Functions, CC: Cellular Component, DNA: Deoxyribonucleic Acid, RNA: Ribonucleic Acid.

| Expression | Category | Term | Count | P-value | Benjamini |
| --- | --- | --- | --- | --- | --- |
| Up-regulated | GOTERM_BP_DIRECT | ubiquitin-dependent protein catabolic process | 13 | 2.3E-5 | 3.5E-2 |
|  | GOTERM_BP_DIRECT | proteasome-mediated ubiquitin-dependent protein catabolic | 12 | 9.1E-5 | 5.1E-2 |
|  | GOTERM_BP_DIRECT | protein stabilization | 12 | 9.8E-5 | 5.1E-2 |
|  | GOTERM_BP_DIRECT | transcription from RNA polymerase II promoter | 11 | 6.4E-4 | 1.5E-1 |
|  | GOTERM_BP_DIRECT | mRNA splicing, via spliceosome | 10 | 6.5E-4 | 1.5E-1 |

|  |  |  |  |  |
| --- | --- | --- | --- | --- |
| GOTERM_BP_DIRECT | behavioral fear response | 5 | 6.8E-4 | 1.5E-1 |
| GOTERM_BP_DIRECT | chromatin remodeling | 13 | 7.1E-4 | 1.5E-1 |
| GOTERM_BP_DIRECT | protein polyubiquitination | 9 | 7.6E-4 | 1.5E-1 |
| GOTERM_BP_DIRECT | cellular response to DNA damage stimulus | 11 | 3.0E-3 | 4.1E-1 |
| GOTERM_BP_DIRECT | peptidyl-serine phosphorylation | 8 | 3.1E-3 | 4.1E-1 |
| GOTERM_MF_DIRECT | RNA binding | 58 | 2.8E-15 | 1.4E-12 |
| GOTERM_MF_DIRECT | protein binding | 200 | 3.5E-11 | 8.6E-9 |
| GOTERM_MF_DIRECT | mRNA binding | 15 | 2.1E-6 | 3.4E-4 |
| GOTERM_MF_DIRECT | DNA binding | 33 | 2.4E-4 | 3.0E-2 |
| GOTERM_MF_DIRECT | transcription coactivator activity | 12 | 4.5E-4 | 4.5E-2 |
| GOTERM_MF_DIRECT | chromatin binding | 16 | 1.0E-3 | 7.1E-2 |
| GOTERM_MF_DIRECT | identical protein binding | 37 | 1.2E-3 | 7.1E-2 |
| GOTERM_MF_DIRECT | damaged DNA binding | 6 | 1.6E-3 | 8.5E-2 |
| GOTERM_MF_DIRECT | double-stranded RNA binding | 6 | 2.0E-3 | 9.9E-2 |
| GOTERM_MF_DIRECT | translation factor activity, RNA binding | 4 | 2.3E-3 | 1.1E-1 |
| GOTERM_CC_DIRECT | nucleus | 122 | 4.4E-14 | 1.7E-11 |
| GOTERM_CC_DIRECT | cytosol | 115 | 9.5E-14 | 1.9E-11 |
| GOTERM_CC_DIRECT | Nucleoplasm | 93 | 1.7E-13 | 2.2E-11 |
| GOTERM_CC_DIRECT | proteasome complex | 9 | 2.4E-7 | 2.4E-5 |
| GOTERM_CC_DIRECT | cytoplasm | 98 | 6.1E-7 | 4.8E-5 |
| GOTERM_CC_DIRECT | extracellular exosome | 47 | 2.8E-5 | 1.8E-3 |
| GOTERM_CC_DIRECT | presynapse | 10 | 1.2E-4 | 7.0E-3 |
| GOTERM_CC_DIRECT | clathrin-coated pit | 6 | 6.7E-4 | 3.0E-2 |
| GOTERM_CC_DIRECT | nuclear membrane | 11 | 7.1E-4 | 3.0E-2 |
| GOTERM_CC_DIRECT | cytoplasmic stress granule | 7 | 7.5E-4 | 3.0E-2 |

**Table 3:**
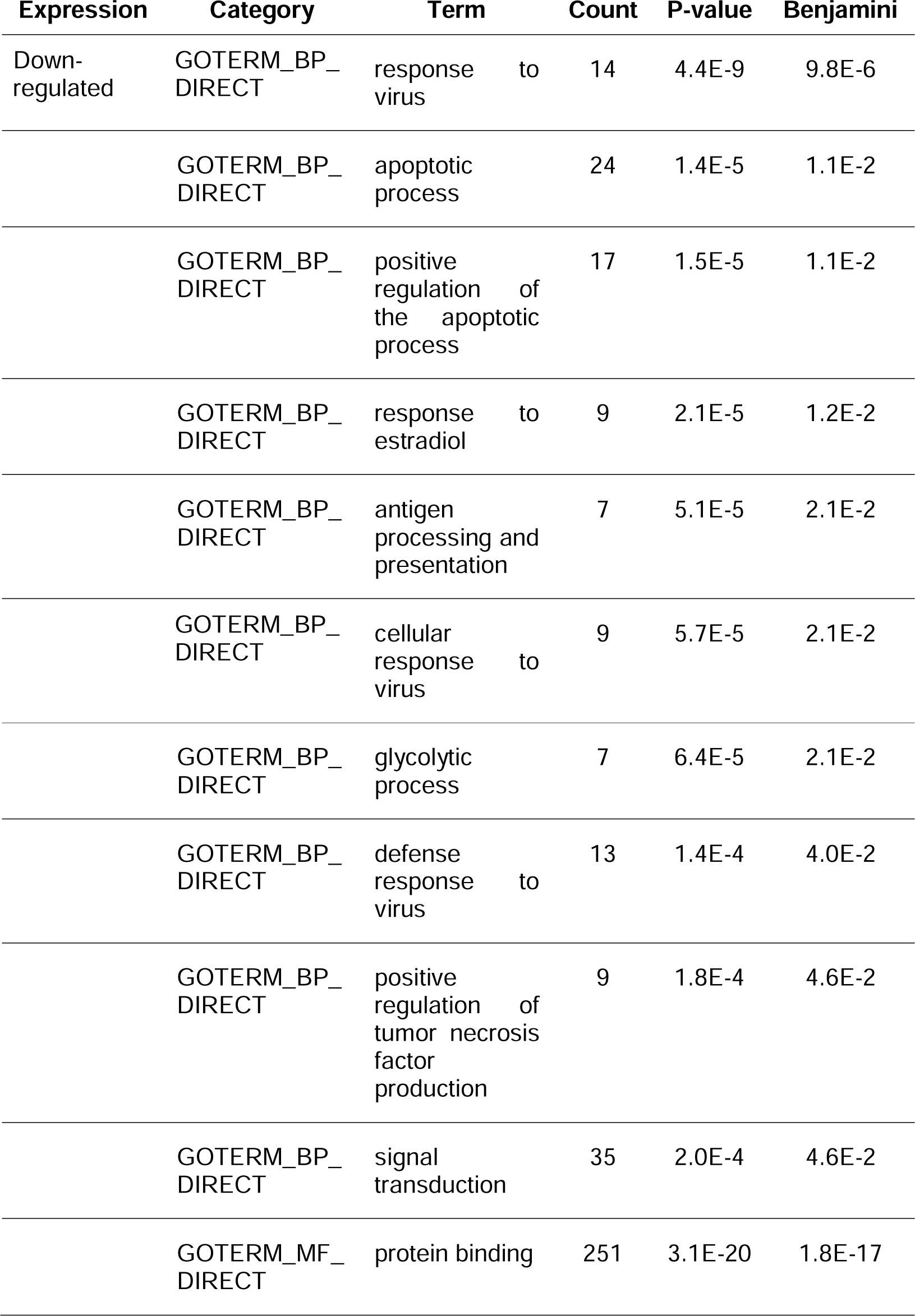
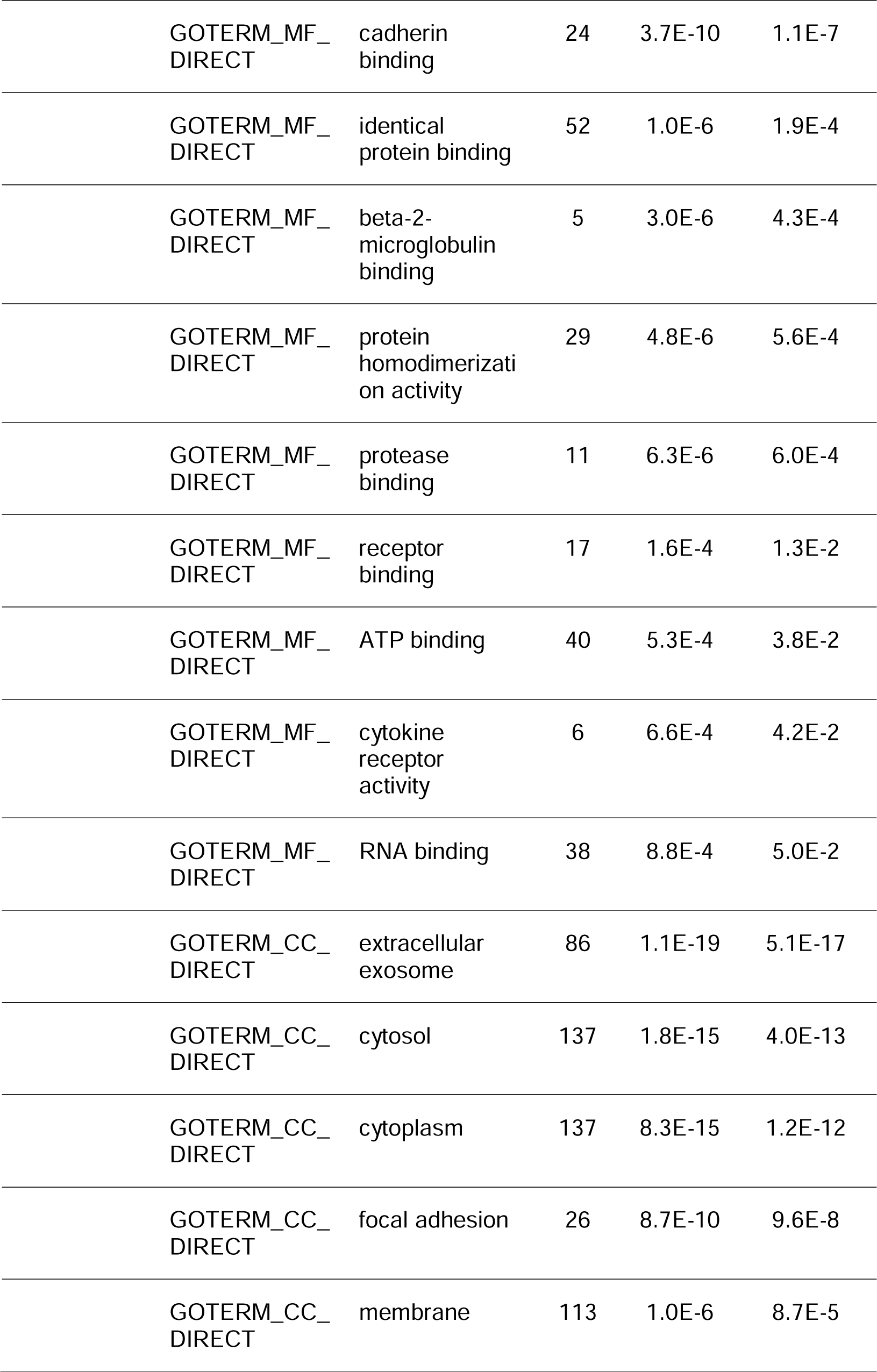
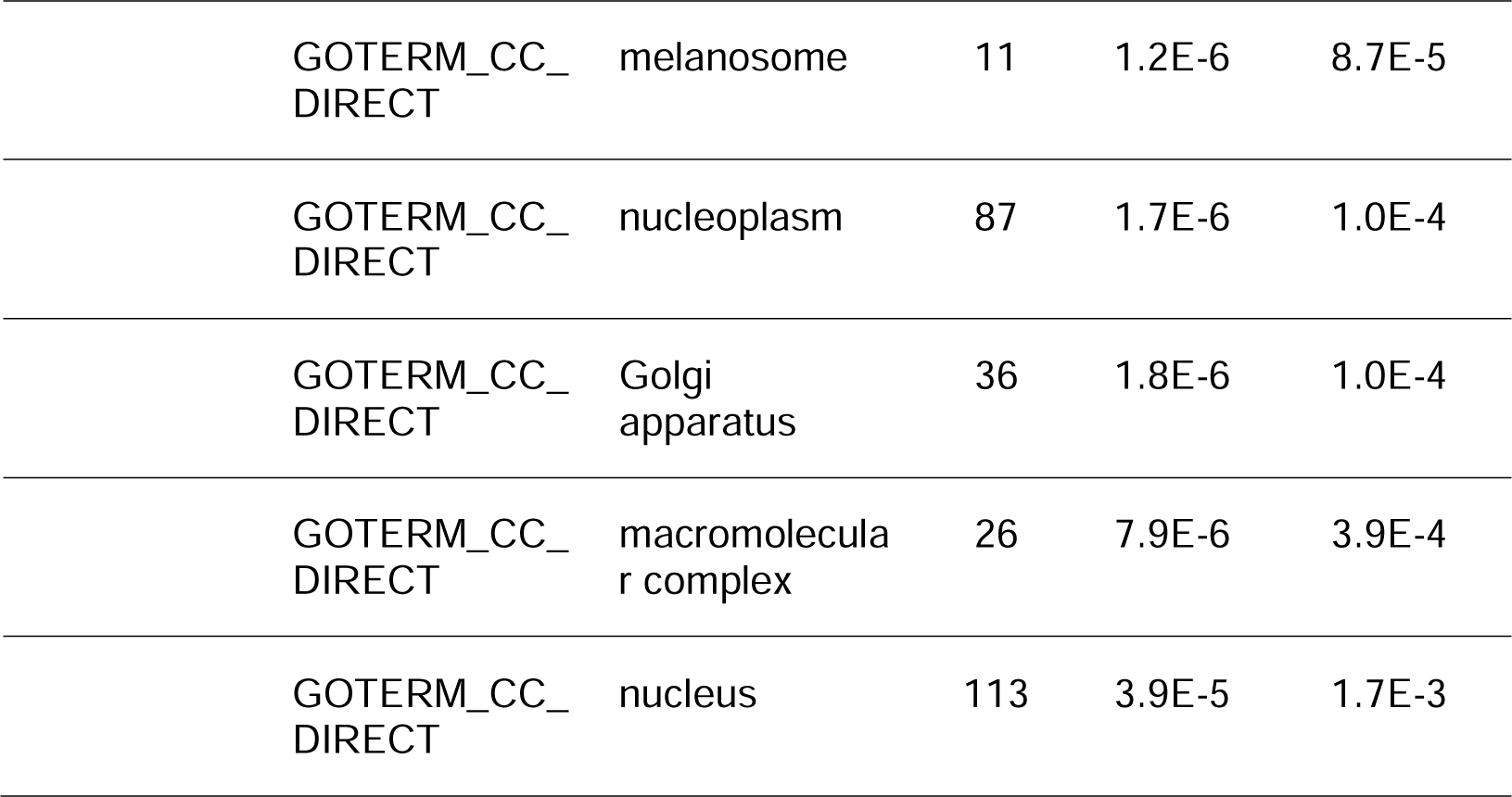
Top 10 Gene Ontology of downregulated DEGs RA. Abbreviations: GOTERM: Gene Ontology Terms, BP: Biological Process, MF: Molecular Functions, CC: Cellular Component, RNA: Ribonucleic Acid.

| Expression | Category | Term |  | Count | P-value | Benjamini |
| --- | --- | --- | --- | --- | --- | --- |
| Down-regulated | GOTERM_BP_DIRECT | response to virus |  | 14 | 4.4E-9 | 9.8E-6 |
|  | GOTERM_BP_DIRECT | apoptotic process |  | 24 | 1.4E-5 | 1.1E-2 |
|  | GOTERM_BP_DIRECT | positive regulation of the apoptotic process |  | 17 | 1.5E-5 | 1.1E-2 |
|  | GOTERM_BP_DIRECT | response to estradiol |  | 9 | 2.1E-5 | 1.2E-2 |
|  | GOTERM_BP_DIRECT | antigen processing and presentation |  | 7 | 5.1E-5 | 2.1E-2 |
|  | GOTERM_BP_DIRECT | cellular response to virus |  | 9 | 5.7E-5 | 2.1E-2 |
|  | GOTERM_BP_DIRECT | glycolytic process |  | 7 | 6.4E-5 | 2.1E-2 |
|  | GOTERM_BP_DIRECT | defense response to virus |  | 13 | 1.4E-4 | 4.0E-2 |
|  | GOTERM_BP_DIRECT | positive regulation of tumor necrosis factor production |  | 9 | 1.8E-4 | 4.6E-2 |
|  | GOTERM_BP_DIRECT | signal transduction |  | 35 | 2.0E-4 | 4.6E-2 |
|  | GOTERM_MF_DIRECT | protein binding |  | 251 | 3.1E-20 | 1.8E-17 |

|  |  |  |  |  |
| --- | --- | --- | --- | --- |
| GOTERM_MF_<br>DIRECT | cadherin<br>binding | 24 | 3.7E-10 | 1.1E-7 |
| GOTERM_MF_<br>DIRECT | identical<br>protein binding | 52 | 1.0E-6 | 1.9E-4 |
| GOTERM_MF_<br>DIRECT | beta-2-microglobulin<br>binding | 5 | 3.0E-6 | 4.3E-4 |
| GOTERM_MF_<br>DIRECT | protein<br>homodimerizati<br>on activity | 29 | 4.8E-6 | 5.6E-4 |
| GOTERM_MF_<br>DIRECT | protease<br>binding | 11 | 6.3E-6 | 6.0E-4 |
| GOTERM_MF_<br>DIRECT | receptor<br>binding | 17 | 1.6E-4 | 1.3E-2 |
| GOTERM_MF_<br>DIRECT | ATP binding | 40 | 5.3E-4 | 3.8E-2 |
| GOTERM_MF_<br>DIRECT | cytokine<br>receptor<br>activity | 6 | 6.6E-4 | 4.2E-2 |
| GOTERM_MF_<br>DIRECT | RNA binding | 38 | 8.8E-4 | 5.0E-2 |
| GOTERM_CC_<br>DIRECT | extracellular<br>exosome | 86 | 1.1E-19 | 5.1E-17 |
| GOTERM_CC_<br>DIRECT | cytosol | 137 | 1.8E-15 | 4.0E-13 |
| GOTERM_CC_<br>DIRECT | cytoplasm | 137 | 8.3E-15 | 1.2E-12 |
| GOTERM_CC_<br>DIRECT | focal adhesion | 26 | 8.7E-10 | 9.6E-8 |
| GOTERM_CC_<br>DIRECT | membrane | 113 | 1.0E-6 | 8.7E-5 |
| GOTERM_CC_<br>DIRECT | melanosome | 11 | 1.2E-6 | 8.7E-5 |
| GOTERM_CC_<br>DIRECT | nucleoplasm | 87 | 1.7E-6 | 1.0E-4 |
| GOTERM_CC_<br>DIRECT | Golgi<br>apparatus | 36 | 1.8E-6 | 1.0E-4 |
| GOTERM_CC_<br>DIRECT | macromolecula<br>r complex | 26 | 7.9E-6 | 3.9E-4 |
| GOTERM_CC_<br>DIRECT | nucleus | 113 | 3.9E-5 | 1.7E-3 |

**Table 4:** Top 10 KEGG pathways of upregulated DEGs associated with RA.

| Expression | Term | Count | P-value | Benjamini |
| --- | --- | --- | --- | --- |
| Up-regulated | Parkinson disease | 18 | 3.8E-6 | 3.7E-4 |
|  | Prion disease | 18 | 5.3E-6 | 3.9E-4 |
|  | Amyotrophic lateral sclerosis | 21 | 5.4E-6 | 3.9E-4 |
|  | Huntington disease | 19 | 6.4E-6 | 3.9E-4 |
|  | Pathways of neurodegeneration multiple diseases | 24 | 8.3E-6 | 4.0E-4 |
|  | Proteasome | 8 | 1.3E-5 | 5.4E-4 |
|  | Alzheimer disease | 19 | 1.3E-4 | 4.4E-3 |
|  | Spinocerebellar ataxia | 11 | 2.0E-4 | 6.0E-3 |
|  | Diabetic cardiomyopathy | 11 | 3.0E-3 | 7.9E-2 |
|  | Pyruvate metabolism | 5 | 9.1E-3 | 2.0E-1 |
|  | Oxidative phosphorylation | 8 | 9.2E-3 | 2.0E-1 |

**Table 5:** Top 10 KEGG pathways of downregulated DEGs associated with RA. Abbreviations: HIF-1: Hypoxia Inducible Factor-1, Th17: T-helper 17.

| Expression | Term | Count | P-value | Benjamini |
| --- | --- | --- | --- | --- |
| Down-regulated | Lysosome | 14 | 6.7E-6 | 1.9E-3 |
|  | HIF-1 signaling pathway | 12 | 2.8E-5 | 3.8E-3 |
|  | Epstein-Barr virus infection | 15 | 1.5E-4 | 1.2E-2 |
|  | Toxoplasmosis | 11 | 1.7E-4 | 1.2E-2 |
|  | Biosynthesis of amino acids | 8 | 1.3E-3 | 7.3E-2 |
|  | Th17 cell differentiation | 9 | 2.7E-3 | 9.2E-2 |
|  | Human papillomavirus infection | 17 | 2.8E-3 | 9.2E-2 |
|  | Inflammatory bowel disease | 7 | 3.1E-3 | 9.2E-2 |
|  | Salmonella infection | 14 | 3.6E-3 | 9.2E-2 |
|  | Glycolysis /Gluconeogenesis | 7 | 3.6E-3 | 9.2E-2 |

On the other hand, the BP for downregulated DEGs includes response to the virus, apoptotic process, positive regulation of the apoptotic process, response to estradiol, and antigen processing and presentation whereas the downregulated DEGs are found highly enriched in protein binding, cadherin binding, identical protein binding, beta-2-microglobulin binding, and protein homodimerization activity in MF. Furthermore, the cellular components of downregulated DEGs are the extracellular exosome, cytosol, cytoplasm, focal adhesion, and membrane. Moreover, the significantly enriched KEGG pathways of upregulated DEGs are Parkinson’s disease, Prion disease, Amyotrophic lateral sclerosis, Huntington’s disease, Pathways of neurodegeneration - multiple diseases while downregulated DEGs exhibited lysosome, HIF-1 signaling pathway, Epstein-Barr virus infection, Toxoplasmosis, and biosynthesis of amino acids.

### Protein-Protein Interaction Network Analysis

The Protein-Protein Interaction (PPI) networks were constructed using the STRING database and consisted of 239 nodes and 892 edges for upregulated DEGs while the downregulated DEGs consisted of 283 nodes and 970 edges. The PPI enrichment p-value is less than 1.0e-16. The PPI network is illustrated in Figure 5 and Figure 6.

**Figure 5:**
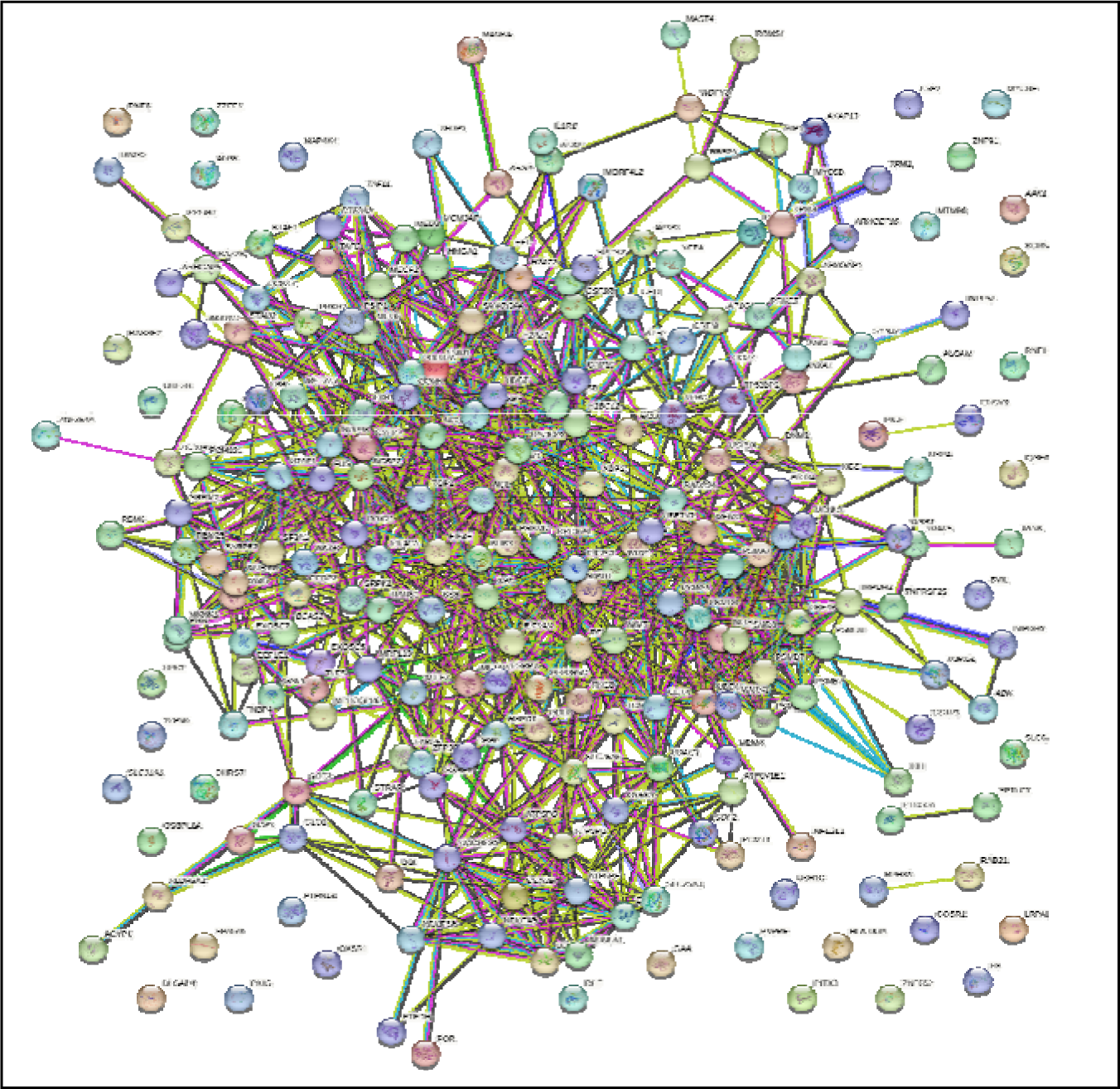
PPI network of upregulated DEGs. 239 nodes and 892 edges.

**Figure 6:**
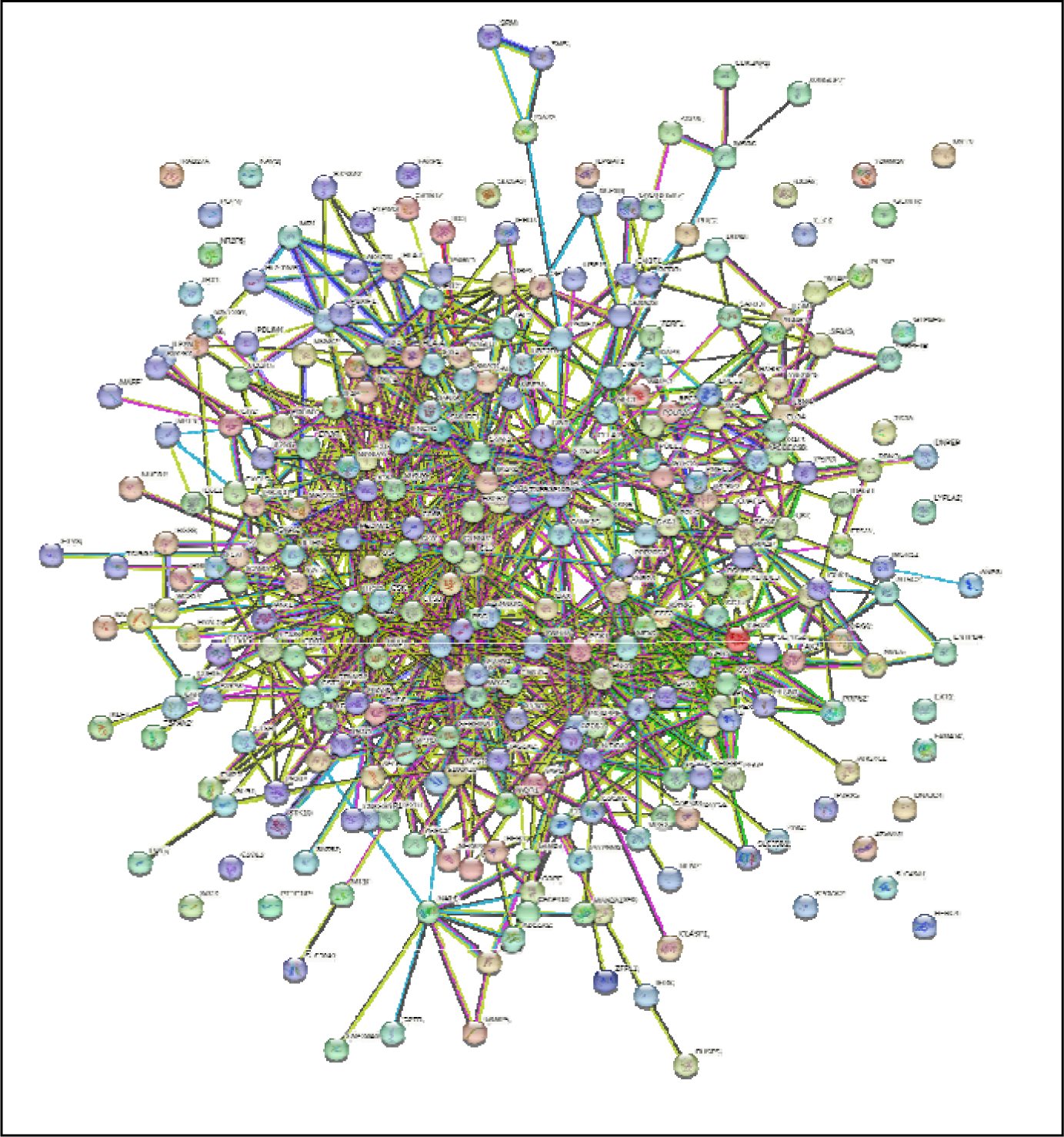
PPI network of downregulated DEGs. 283 nodes and 970 edges.

### Identification of Hub Genes

The 10 topmost hub genes associated with RA were identified using Cytohubba plugged in Cytoscape v 3.9.1. The top 10 upregulated hub genes include RPS27A, UBB, UBC, UBA52, PSMD4, PSMD1, PSMD7, PSMB7, PSMD8 and PSMA7 whereas ACTB, TP53, AKT1, GAPDH, CTNNB1, EGFR, TNF, IL6, MYC and ANXA5 are for downregulated DEGs as shown in Figure 7. The results revealed that RPS27A and TP53 hub genes are closely associated with RA for upregulated and downregulated hub genes respectively as they are the first-rank genes in the top 10 and play key roles in the pathogenesis of RA.

**Figure 7:**
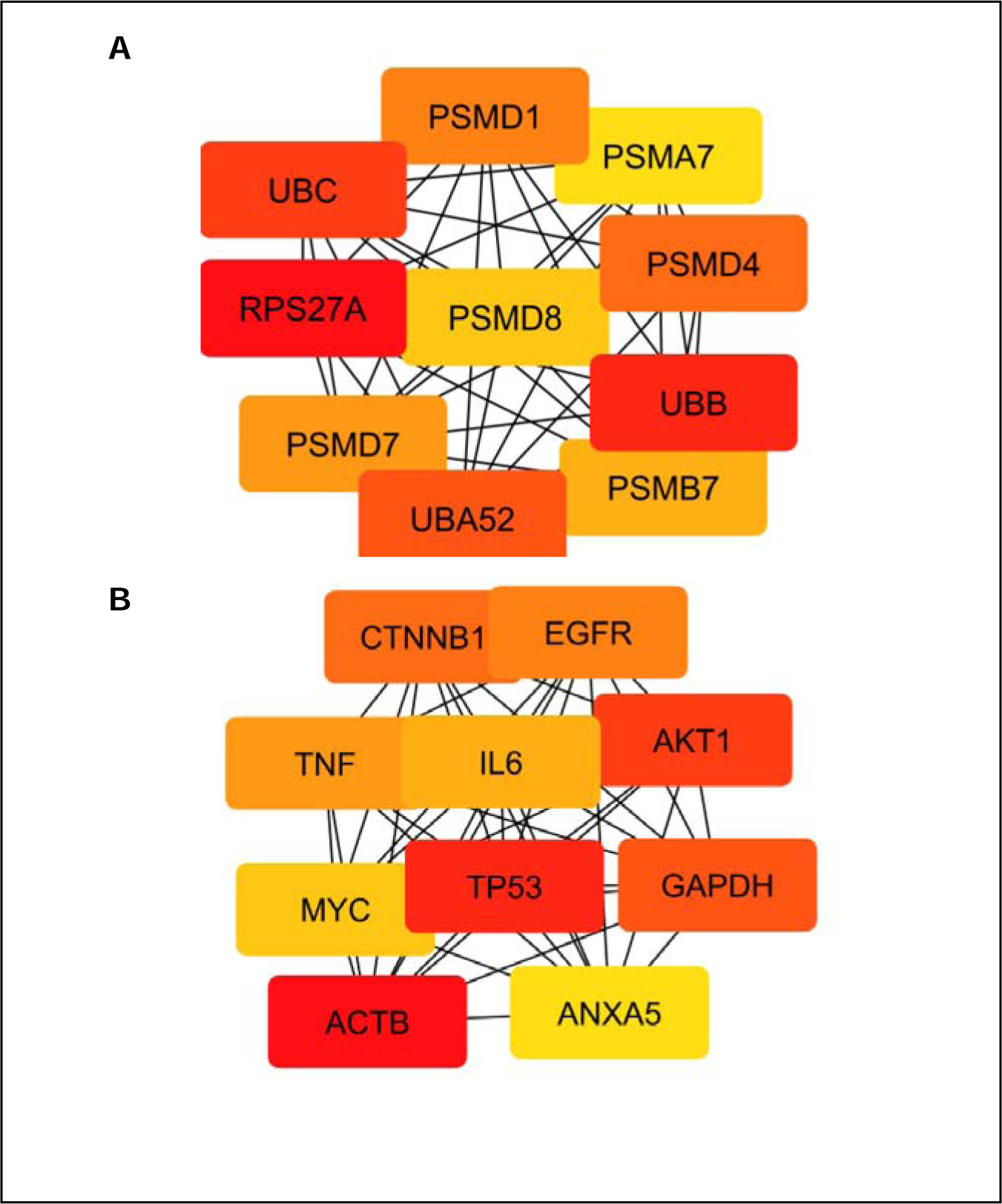
Identification of the top 10 hub genes associated with RA. **(A)** Top 10 upregulated hub genes. **(B)** Top 10 downregulated hub genes.

### MiRNA Network Analysis

The miRNA networks for upregulated and downregulated hub genes are shown in Figure 8. The results depict that 105 edges connect 29 nodes of miRNA, and 10 nodes of upregulated hub genes, and hsa-mir-23b-3p was found to have the highest degree of association with the top 10 upregulated hub genes with a value of 6. On the other hand, 480 edges connect 114 nodes of miRNA and 10 nodes of downregulated hub genes. hsa-mir-34a-5p and hsa-mir-155-5p exhibited the highest degree of association, a value of 10 with the top 10 downregulated hub genes.

**Figure 8:**
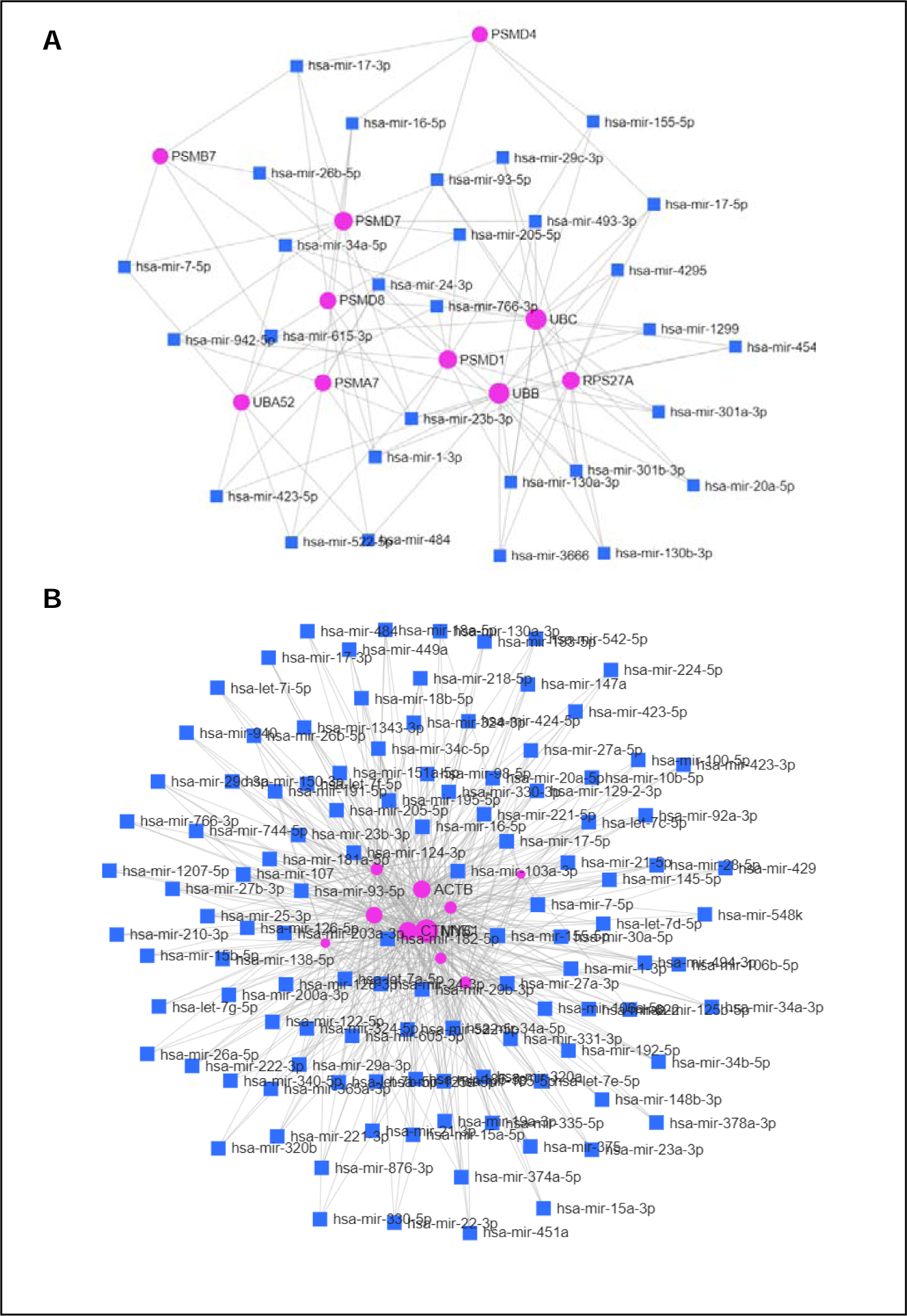
miRNA networks of hub genes associated with RA. Pink circles represent the downregulated hub genes associated with RA. The blue square indicates the miRNA targeting hub genes. **(A)** mi-RNA network of upregulated hub genes. **(B)** miRNA networks of downregulated hub genes.

### Identification of Immune Response

The GO terms and KEGG pathways associated with the RA immune response of RPS27A and TP53 are obtained from InnateDB (Table 6 and Table 7). Both of these hub genes play key roles in triggering the immune response during the pathogenesis of RA.

**Table 6:** GO terms and KEGG pathways of RPS27A related to immune response of Rheumatoid Arthritis using InnateDB. Abbreviations: GO: Gene Ontology.

| Biological Process | Molecular Function | Cellular Component | KEGG |
| --- | --- | --- | --- |
| GO:0045087<br>innate immune response | GO:0003735<br>structural constituent of ribosome | GO:0005622<br>intracellular | hsa03010<br>Ribosome pathway |
| GO:0019221<br>cytokine-mediated signaling pathway | GO:0005515<br>protein binding | GO:0005829<br>cytosol |  |
| GO:0007249<br>I-kappaB kinase/NF-kappaB signaling | GO:0046872<br>metal ion binding | GO:0005840<br>ribosome |  |

**Table 7:** GO terms and KEGG pathways of TP53 related to immune response of Rheumatoid Arthritis using InnateDB. Abbreviations: GO: Gene Ontology.

| **Biological Process** | **Molecular Function** | **Cellular Component** | **KEGG** |
| --- | --- | --- | --- |
| GO:0045087<br>innate immune response | GO:0002039<br>p53 binding | GO:0000785<br>chromatin | hsa04310<br>Wnt signaling pathway |
| GO:0002326<br>B cell lineage commitment | GO:0031625<br>ubiquitin protein ligase binding | GO:0000790<br>nuclear chromatin | hsa04210<br>Apoptosis pathway |
| GO:0002309<br>T cell proliferation involved in immune response | GO:0003684<br>damaged DNA binding | GO:0005634<br>nucleus | hsa04010<br>MAPK signaling pathway pathway |

### Retrieval of *Celastrus Paniculatus* Compounds

A total of 20 compounds of *C.paniculatus* were obtained from the IMPPAT database. Precise information about the plant including kingdom, family, common name, synonym name, and system of medicine was also retrieved from the IMPPAT database as shown in Table 8. Based on the table, *C.paniculatus* belongs to the Celastraceae family, commonly known as the Black Oil plant. It has been used in Indian Traditional Medicines such as Ayurved, Siddha, Sowa Rigpa and Unani.

**Table 8:**
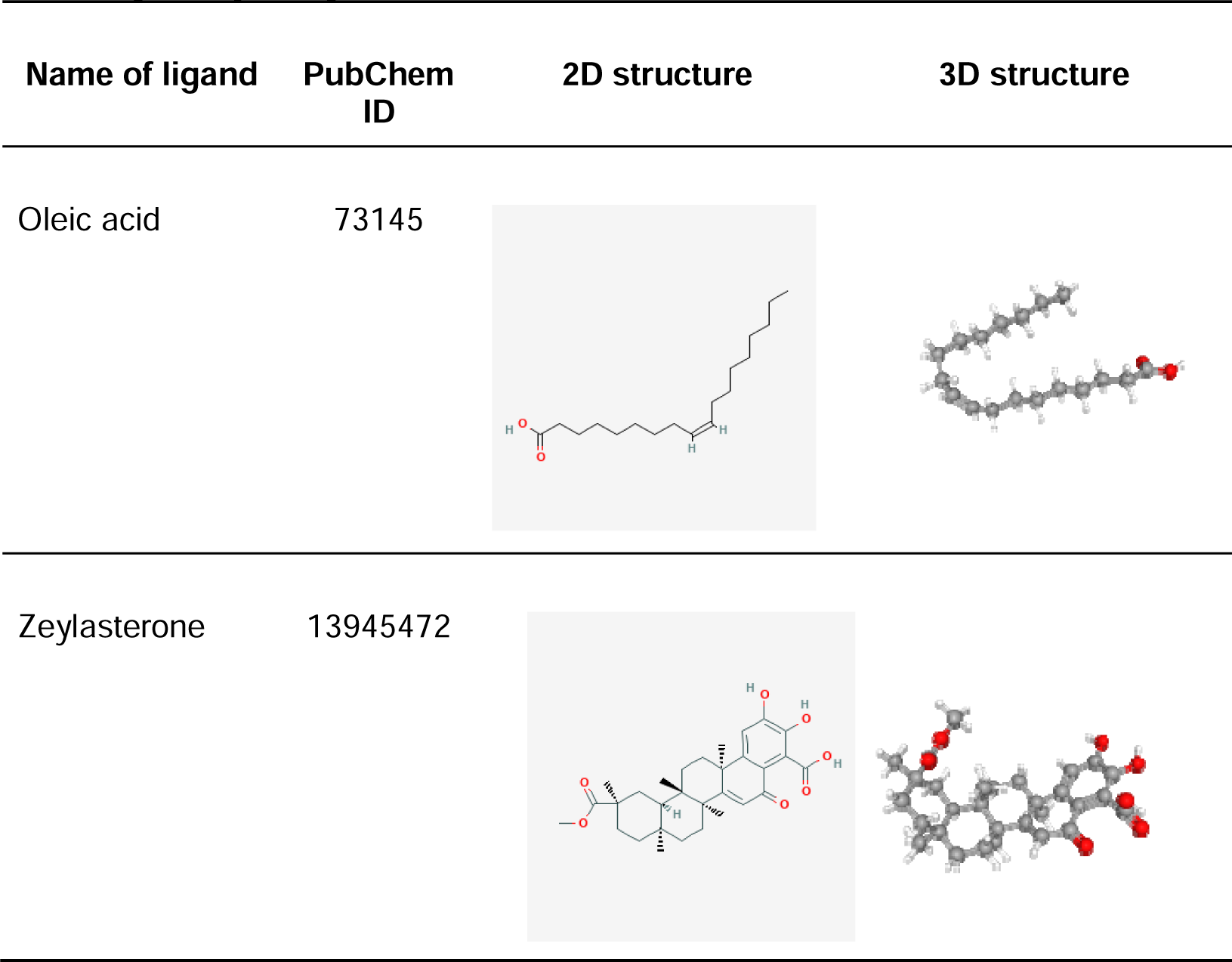
Two-dimensional and three-dimensional structure of ligands with strongest binding affinity. Image obtained from : PubChem, 2004.

| Name of ligand | PubChem ID | 2D structure | 3D structure |
| --- | --- | --- | --- |
| Oleic acid | 73145 |  |  |
| Zeylasterone | 13945472 |  |  |

### Autodocking

RPS27A and TP53 targeted proteins were docked with 20 compounds of C.paniculatus using Autodock Vina v1.5.7. The active binding sites of the targeted proteins were analyzed and generated into a grid file. The dimensions of the grid file reported that x = 19.171, y = 2.013, and z = 23.725 for RPS27A whereas x = 4.353, y = 12.176, and z = -23.731 for TP53.

The docking results revealed that oleic acid exhibited the most effective binding interactions with RPS27A compared to other ligands as evidenced by its binding affinity, -6.4kcal/mol. On the other hand, zeylasterone showed the strongest binding affinity with TP53 with a value of -7.8kcal/mol among the other ligands. The two-dimensional and three-dimensional structures of the ligands and targeted proteins with strong binding affinity are demonstrated in Table 9 and Table 10 respectively. The comparison list of ligands and their respective binding affinities are shown in Table 11.

**Table 9:**
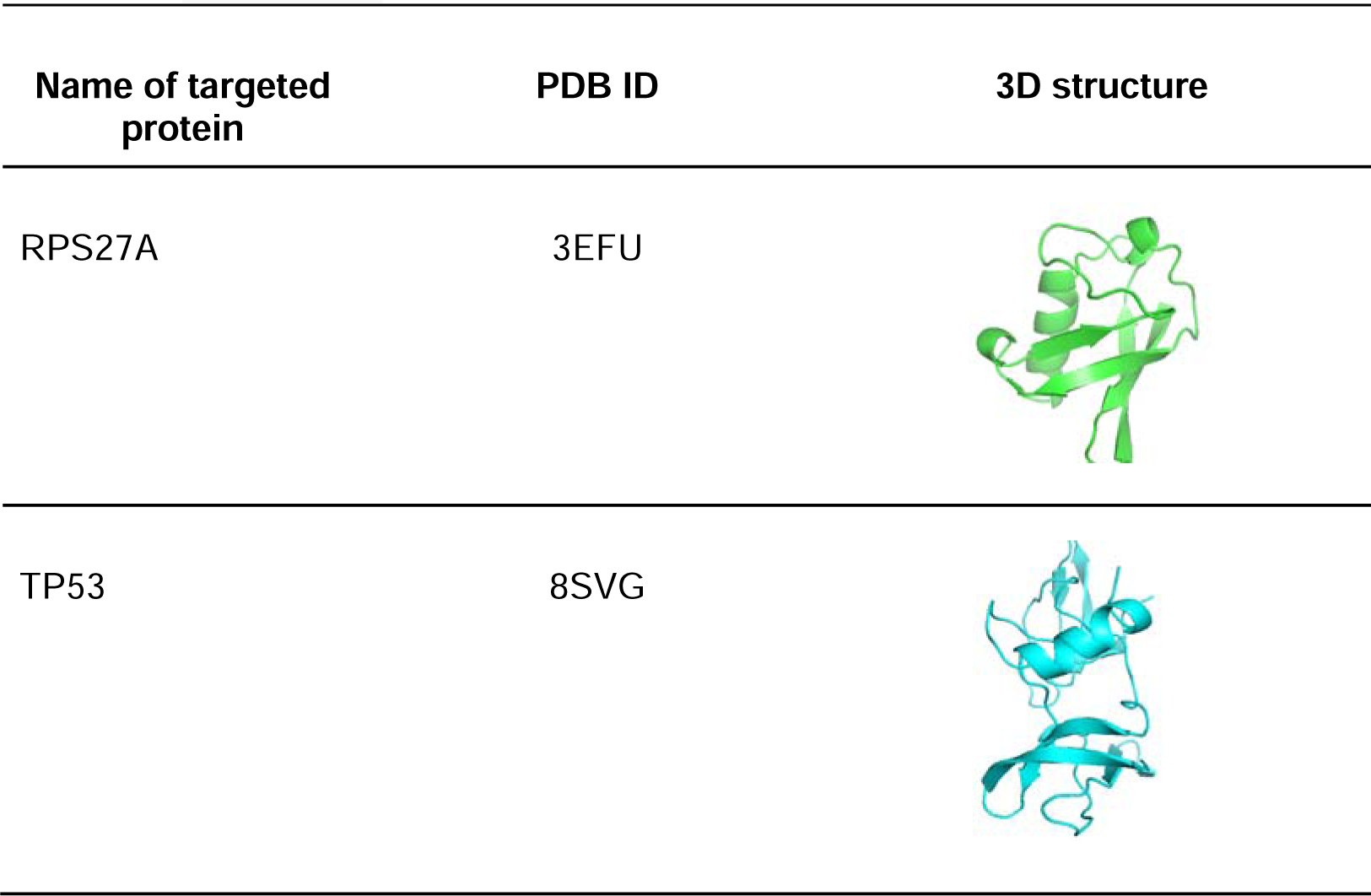
Two-dimensional and three-dimensional structure of targeted proteins vizualized using Pymol. Abbreviations: RPS27A: Ribosomal Protein S27a, TP53: Tumour Protein P53.

| Name of targeted protein | PDB ID | 3D structure |
| --- | --- | --- |
| RPS27A | 3EFU |  |
| TP53 | 8SVG |  |

**Table 10:** Binding affinities of ligands with RPS27A and TP53 targeted proteins. Abbreviations: RPS27A: Ribosomal Protein S27a, TP53: Tumour Protein P53.

| LIGANDS | BINDING AFFINITIES (Kcal/mol) |  |
| --- | --- | --- |
|  | RPS27A | TP53 |
| Benzoic acid | -4.2 | -4.8 |
| Pristimerin | -6.3 | -7.4 |
| Zeylasteral | -6.0 | -7.6 |
| Celastrol | -6.3 | -7.6 |
| Zeylasterone | -6.3 | -7.8 |
| Stearic acid | -3.4 | -4.0 |
| 1-Triacontanol | -2.8 | -4.1 |
| Tetracosanoic acid | -3.3 | -4.8 |
| (1S,3S,8S,9R,12S,14R,16S)-16-hydroxy-5,14-dimethyl-5-azatetracyclo[7.7.0.0.1,12.0.3,8]hexadecan-10-one | -5.3 | -5.8 |
| 1-Tetracosanol | -3.0 | -4.2 |
| Celapanine | -5.1 | -6.4 |
| Paniculatadiol | -3.0 | -4.1 |
| Palmitic acid | -3.8 | -4.4 |
| Malkangunin | -5.4 | -6.5 |
| Malkanguniol | -4.9 | -6.3 |
| Oleic acid | -6.4 | -7.6 |
| beta-Amyrin | -3.7 | -4.2 |
| Linolenic acid | -5.7 | -6.4 |
| beta-Sitosterol | -3.9 | -4.3 |
| Linoleic acid | -3.6 | -4.5 |

**Table 11:** Drug-likeness properties of Oleic acid and Zeylasterone compounds of C.paniculatus. TPSA: Topological Polar Surface Area. iLOG P: implicit logarithm P, ESOL LOG S: Estimated Solubility Logarithm of Solubility.

| Compounds | Drug-likeness |  |  |  |  |  |  |
| --- | --- | --- | --- | --- | --- | --- | --- |
|  | Molecular Weight<br>(g/mol) | No. H-bond<br>acceptors | No. H-bond donors | TPSA (Å²) | iLOGP | ESOL LOG S | Lipinski's Rule |
| Oleic acid | 282.46 | 2 | 1 | 37.30 | 4.27 | -5.41 | YES |
| Zeylasterone | 510.62 | 7 | 3 | 121.13 | 3.49 | -7.09 | YES |

The binding sites and interactions involved in docking were viewed and analyzed using Pymol and Discovery Studio. The analysis revealed that two polar residues namely, His 68 (Histidine 68) and Phe 4 (Phenylalanine 4) were involved in pi-alkyl interactions between oleic acid and RPS27A targeted protein (Figure 9). In addition, His 233 (Histidine 233), Asn 200 (Asparagine 200), Glu 198 (Glutamic acid 198), and Ser 227 (Serine 227) participate in hydrogen interactions whereas Ile 232 (Isoleucine 232) and Cys 229 (Cysteine 229) are involved in alkyl interactions between zeylasterone and TP53 (Figure 10).

**Figure 9:**
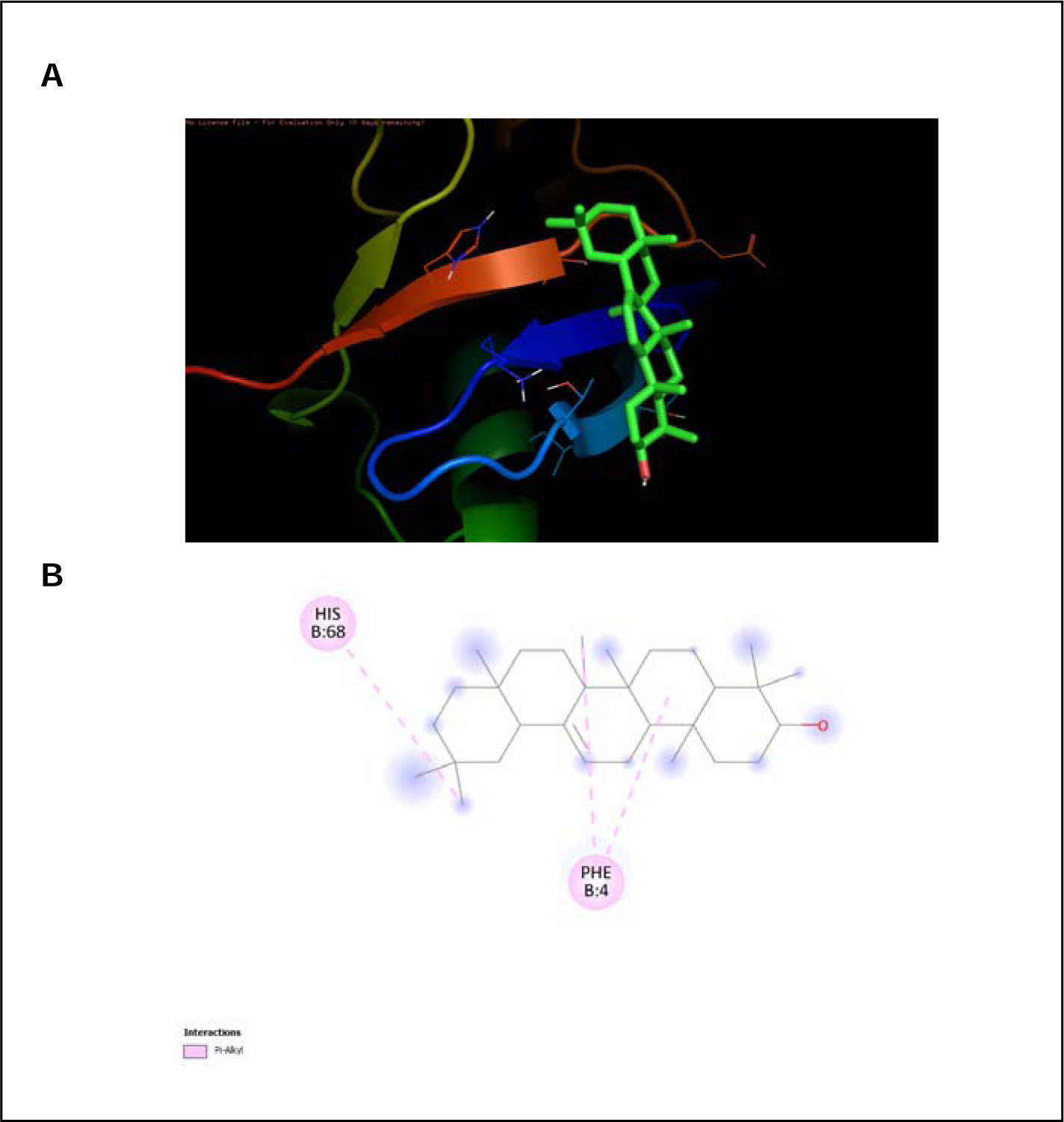
Autodocking result of Oleic acid and RPS27A. **(A)** Three-dimensional binding of Oleic acid at the active site of RPS27A. **(B)**Two-dimensional structure depicting interactions with amino acid residues involved in hydrophobic interaction. RPS27A: Ribosomal Protein S27a, HIS 68: Histidine 68, PHE 4: Phenylalanine 4.

**Figure 10:**
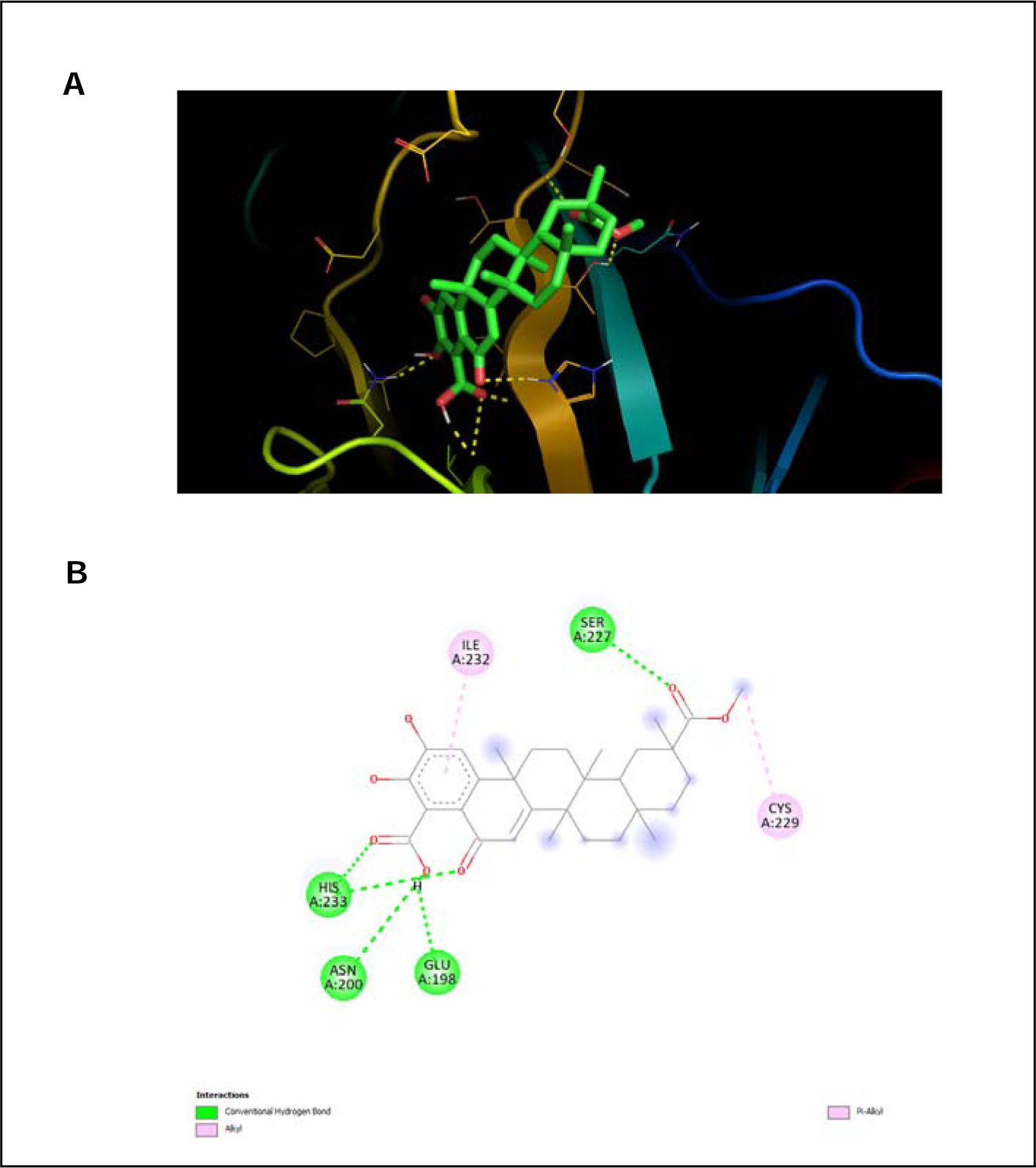
Autodocking result of Zeylasterone and TP53. **(A)** Three-dimensional binding of Zeylasterone at the active site of TP53. **(B)** Two-dimensional structure depicting interactions with amino acid residues involved in hydrogen and hydrophobic interactions.TP53: Tumor Protein P53, His 233: Histidine 233, Asn 200: Asparagine 200, Glu 198: Glutamic acid 198, Ser 227: Serine 227, Ile 232: Isoleucine 232, Cys 229: Cysteine 229.

### Drug-likeness and Pre-Admet Analysis

The drug-likeness helps to predict the drug’s nature and oral bioavailability which complies with Lipinski’s rules while the PreADMET analysis is a valuable analysis to predict features of the compounds such as the toxicological and pharmacological properties specifically during the pre-clinical phase (Ahmad et al., 2021)(Noor et al., 2019). This approach contributes to the optimization of medication and prevents late-stage failures in drug discovery (Sharma et al., 2023). The drug-likeness property and major parameters : (CaCo-2 permeability, Pgp-inhibitor, Pgp-substrate, Human Intestinal Absorption (HIA), Plasma Protein Binding (PPB), Blood-brain barrier (BBB), Cytochrome P450 (CYP) inhibition) of ADME properties of oleic acid and zeylasterone compounds are tabulated in Table 12, Table 13, Table 14. In addition, the toxicity test was also performed to predict the hERG blockers, AMES toxicity, rat oral acute toxicity, and carcinogenicity of the compounds (Table 15).

**Table 12:** ADME properties of Oleic acid and Zeylasterone compounds of *C.paniculatus.* CaCo-2: colon adenocarcinoma cell lines, Pgp: P-glycoprotein, PPB: Plasma Protein Binding, BBB: Blood-brain barrier.

| Compounds | PreADMET_ADME |  |  |  |  |  |
| --- | --- | --- | --- | --- | --- | --- |
|  | CaCo-2 Permeability | Pgp-inhibitor | Pgp-substrate | Human Intestinal Absorption | PPB (%) | BBB (cm/s) |
| Oleic acid | -4.922 | NO | NO | 0.007 | 98.76 | 0.101 |
| Zeylasterone | -5.64 | NO | NO | 0.02 | 98.13 | 0.046 |

**Table 13:** Cytochrome P450 inhibition properties of Oleic acid and Zeylasterone compounds of *C.paniculatus.* CYP: Cytochrome P450.

| Compounds | PreADMET_CYP INHIBITION |  |  |  |  |
| --- | --- | --- | --- | --- | --- |
|  | CYP1A2 | CYP2C19 | CYP2C9 | CYP2D6 | CYP3A4 |
| Oleic acid | 0.32 | 0.184 | 0.256 | 0.007 | 0.054 |
| Zeylasterone | 0.013 | 0.022 | 0.156 | 0.049 | 0.317 |

**Table 14:** Toxicity properties of Oleic acid and Zeylasterone compounds of *C.paniculatus.* hERG: human-ether-a-go-go-related gene, AMES: Ames Mutagenicity Assay.

| Compounds | PreADMET_Toxicity |  |  |  |
| --- | --- | --- | --- | --- |
|  | hERG Blockers | AMES toxicity | Rat Oral Acute Toxicity | Carcinogenicity |
| Oleic acid | 0.038 | 0.004 | 0.016 | 0.084 |
| Zeylasteral | 0.042 | 0.424 | 0.042 | 0.807 |

As presented in Table 12, oleic acid complies with all of Lipinski’s rules. Moreover, as illustrated in Table 13, oleic acid is CaCo-2 permeable with a value of - 4.942, non-Pgp-inhibitor, non-Pgp-substrate, and high HIA with a value of 0.013 which shows its rapid intestinal absorption. Moreover, it is found that it has high plasma protein binding with 98.76%, and does not penetrate the blood-brain barrier indicating that it is poorly distributed to the brain.

On the other hand, zeylasterone also complies with Lipinski’s rules. It is non-permeable to CaCo-2 with a value of -5.64, non-Pgp-inhibitor, non-Pgp-substrate, and has high HIA as evidenced by its value of 0.007. However, it has a high plasma protein binding with a value of 97.11% and it also does not penetrate the blood-brain barrier. In addition, as presented in Table 14, both oleic acid and zeylasterone are found to be non-inhibitors of all cytochrome P450 family.

Other important aspects of PreADMET are hERG blocker, AMES test, rat oral acute toxicity, and carcinogenicity. The results revealed that the oleic acid is inactive in inhibition of the hERG, AMES negative, deficient rat oral acute toxic, and non-carcinogen. Similarly, zeylasterone is also inactive in blocking the hERG, AMES negative, very low rat oral acute toxicity, and non-carcinogen.

## DISCUSSION

RA is an acute autoimmune disease significantly affecting the diarthrodial joints of the synovium leading to loss of mobility and physical functions (Wu et al., 2021). It has become one of the common factors of disabilities in RA patients (Wu et al., 2021). Since the pathogenesis of RA is complex and still unclear, RA has become a topic of broad and sparked the interest of researchers in the field of immunity (Luo et al., 2023; Wu et al., 2021). Identification of potential hub genes and immune responses associated with RA can provide fundamental insights to unfold the pathogenesis of RA (Luo et al., 2023). Therefore, we analysed the 3 chosen microarray datasets to screen DEGs associated with RA and identified the intersected DEGs among the three datasets which were 243 for upregulated DEGs and 285 for downregulated DEGs. The DEGs were subjected to further GO and KEGG analysis to gain insights into biological functions and signalling pathways which are significantly enriched by the DEGs. The GO and KEGG pathways were visualised in a barplot using -log10(p-value) which is a transformed logarithm expression of P-value. Hence, the higher the -log10(p-value), the more significant the result. Based on Table 4.2.1, it is found that the upregulated DEGs are highly enriched in ubiquitin-dependent protein catabolic processes for BP, RNA binding for MF, and nucleus for CC. Based on Table 4.2.2, the downregulated DEGs are enriched in response to virus for BP, protein binding for MF, and extracellular exosomes for CC.

Abnormal gene expression is closely associated with the onset and progression of RA. In this study, the top 10 potential hub genes associated with RA are discovered, namely RPS27A, UBB, UBC, UBA52, PSMD4, PSMD1, PSMD7, PSMB7, PSMD8, PSMA7 for top 10 upregulated hub genes while ACTB, TP53, AKT1, GAPDH, CTNNB1, EGFR, TNF, IL6, MYC, ANXA5 are the 10 top-most downregulated hub genes. These hub genes are potential diagnostic targets of RA.

Ribosomal Protein S27a (RPS27A), Ubiquitin B (UBB), Ubiquitin C (UBC), and Ubiquitin A-52 Residue Ribosomal Protein Fusion Product 1 (UBA52) are the protein-coding genes from the Ubiquitin family that encode the same protein which is Ubiquitin (Luo et al., 2023). It is a conserved protein and plays an essential role in degradation by targeting cellular proteins, maintaining the structure of chromatin, gene expression regulation, DNA repair, and cell signalling pathways (Luo et al., 2023). The Ubiquitin protein activates the Ubiquitin Proteasomal Pathway (UPP) which carries out cell cycle progression, cell division, apoptosis, and immune response regulation (Behl et al., 2020). Hence, studies have confirmed that when alteration occurs to these genes, it can disrupt the molecular functions and cause high production of the Ubiquitin protein leading to chronic joint damage due to its dysregulation (Behl et al., 2020).

Moreover, the Proteasome 26S Subunit Ubiquitin Receptor Non-ATPase 4 (PSMD4), Proteasome 26S Subunit Non-ATPase 1 (PSMD1), Proteasome 26S Subunit Non-ATPase 7 (PSMD7), Proteasome 20S Subunit Beta 7 (PSMB7), Proteasome 26S Subunit Non-ATPase 8 (PSMD8), and Proteasome 20S Subunit Non-ATPase Alpha 7 (PSMA7) are protein-coding genes from the Proteosome family. These genes maintain protein homeostasis by discarding misfolded or damaged proteins which can disrupt cellular functions and mediate the ubiquitin-independent protein degradation which is essential in several pathways such as the production of subset MHC I (Zhang et al., 2021). Previous studies demonstrated that PSMA7 and the genes from the same family are potential targets for the development of inflammatory diseases as they can interact with the proteins involved in tumour initiation and progression (Behl et al., 2020). Thus, it has a high chance of becoming the biomarker of RA.

Furthermore, the 10 topmost downregulated hub genes associated with RA are shown in Figure 4.4.1. The results demonstrate that AKT Serine / Threonine Kinase 1 (AKT1) and Epidermal Growth Factor Receptor (EGFR) are protein kinase family members. Both of these genes are responsible for various signalling pathways specifically involved in the binding of membrane-bound ligands. AKT1 and EGFR genes are key players in various types of cancer. Besides, Jang et al. (2022) stated that AKT protein is involved in the pathogenesis of RA by triggering immune cell activation and joint inflammation because the AKT protein signalling pathway is essential for tumour cell survival. In addition, EGFR is associated with the regulation of joint damage in RA, suggesting its potential role in the onset of RA (Jang et al., 2022).

Additionally, Interleukin 6 (IL6) is a gene that encodes cytokines for inflammation and differentiation of B cells. It encodes protein at the site of chronic inflammation causing inflammation-associated diseases such as RA (Favalli, 2020). Studies have unravelled that IL6 regulates innate and adaptive immunity causing it to play a role in cellular processes such as activating immune cells such as T-cells, B-cells and interacting with immune cells leading to RA development and progression (Favalli, 2020; Jarlborg & Gabay, 2022). Ogata et al. (2019) state that dysregulated IL6 has the potential to cause joint inflammation, stiffness, and cartilage degradation.

Moreover, the Tumour Protein P53 (TP53) gene is also one of the hub genes of RA. It encodes the p53 tumour suppressor protein which suppresses the inflammation in RA (Taghadosi et al., 2021). P53 also regulates the signalling pathway of inflammation, induces cytokines, and modulates immune responses (Taghadosi et al., 2021). Studies have revealed that somatic mutations or low expression in TP53 occur in synovial tissue of RA patients causing inflammatory responses and cytokine production such as IL17 and TNF-alpha (Firestein et al., 1997; Yamanishi et al., 2002). Hence, the dysregulation of TP53 can cause p53 protein to be dysfunctional, leading to not being able to regulate inflammation pathways and eventually causing RA (Taghadosi et al., 2021).

The Tumour Necrosis Factor (TNF) is one of the top 10 downregulated hub genes that is implicated in the pathogenesis of RA. It is from the tumour necrosis factor family which is a proinflammatory cytokine. Hence, it plays a major role in triggering inflammatory responses in RA (Farrugia & Baron, 2016). Previous studies have revealed that abnormal expression of TNF can cause acute inflammation, systemic manifestation, and joint damage (Farrugia & Baron, 2016).

Moreover, the Actin Beta (ACTB) plays a key role in the pathogenic activation of synovial fibroblasts (SFs) (Wu et al., 2022). This activation causes aberrant changes in morphology and biochemical behaviour developing chronic destructive inflammation in joints (Wu et al., 2022). In addition, Glyceraldehyde-3-Phosphate Dehydrogenase (GAPDH) is a gene that regulates aerobic glycolysis and immunomodulation (Harshan et al., 2022). Abnormal expression of it in hypoxic synovial tissue can cause inflammation in arthritis patients (Harshan et al., 2022). On the other hand, MYC Proto-Oncogene (MYC) is also one of the top 10 downregulated hub genes. It encodes c-MYC protein which regulates the growth and invasiveness of rheumatoid arthritis synovial fibroblast (RASF) and inhibits apoptosis of RASF leading to promotion of inflammation (Zacarías-Fluck et al., 2024). It also activates immune cells such as T-cells which significantly contribute to the pathogenesis of RA (Zacarías-Fluck et al., 2024). Furthermore, Annexin A5 (ANXA5) is one of the hub genes that has been extensively studied for its role in RA (Baek et al., 2017). It is implicated in the regulation of apoptosis and inflammation in RA (Baek et al., 2017). However, the aberrant expression of it can cause dysregulation of apoptosis and inflammation leading to chronic joint inflammatory responses (Baek et al., 2017).

Besides, the top 10 upregulated and downregulated hub genes explained above were used to construct miRNA-hub genes network. miRNA-hub genes network analysis is essential to comprehend the intricate role of miRNAs in regulating the transcripts, biological pathways, immune differentiation, immune responses and interactions of cells involved in RA such as immune cells, osteoblasts, osteoclasts, and fibroblast-like synoviocytes (FLS) (C. Chang et al., 2022; Peng et al., 2023; Yang et al., 2022). Studies have proved that miRNA is directly involved in regulating inflammatory cytokines, promoting immune cell differentiation into cells like Th17, making it a potential target for gene therapy (Chang et al., 2022). Hence, targeting miRNA provides promising avenues for the discovery of novel treatment approaches, and biomarkers for early diagnosis of RA.

The miRNA network targeting the upregulated and downregulated hub genes of RA is illustrated in Figure 4.5.1. The result reveals that hsa-mir-23b-3p regulates gene expression in RA as it has the highest degree of association with a value of 6 compared to all the other miRNAs in upregulated hub genes. Previous studies conducted by Han et al. (2023) showed that mir-23b-3p acts as a joint inflammation key suppressor and inhibits the progression of RA. It balances the resistant cells such as fibroblast-like synoviocytes (FLS) lining in the inflamed synovium by influencing the NF-kB signalling pathway (Han et al., 2023). Additionally, mir-23b-3p can reduce the secretion of proinflammatory cytokines such as TNF, IL6, and IL-beta which can diminish autoimmune inflammation (Han et al., 2023). In contrast, it was found that hsa-mir-23b-3p is overexpressed in FLS and high plasma levels of hsa-mir-23b-3p in RA patients compared to normal healthy people (Liu et al., 2019). A high plasma level of hsa-mir-23b-3p increases the activity of RA markers such as erythrocyte sedimentation rate (ESR) and C-reactive protein (CRP) (Liu et al., 2019). This proves that greater expression of hsa-mir-23b-3p can lead to proliferation of FLS which is a cell responsible for synovial inflammation and joint damage, and high secretion of cytokines. Hence, mir-23b-3p can be a potential RA novel biomarker and therapeutic target, contributing promising approaches for therapies such as miRNA mimics, and antagonists to control the miRNA expression (Peng et al., 2023; Rezaeepoor et al., 2020; Zhang et al., 2023).

Furthermore, the hsa-mir-34a-5p and hsa-mir-155-5p are found to have the highest degree of association with a value of 10 compared to other miRNAs for downregulated hub genes. The hsa-mir-34a-5p helps in regulating gene expression for RA. Previous research has identified that mir-34a-5p inhibits the expression of 3’UTR of SIRT1, leading to high acetylation of p53 and promoting apoptosis of FLS in RA (Zhang et al., 2023). However, the result depicted the low expression of mir-34a-5p which leads to low acetylation of p53 and hyperproliferation of FLS in RA patients. On the other hand, the hsa-mir-155-5p helps to regulate the activation and responses of B cells in RA (Renman et al., 2020). The dysregulation of hsa-mir-155-5p can cause cytokine production such as IL-6, IL-17, and TNF-alpha (Rezaeepoor et al., 2020). Thus, studies suggested that hsa-mir-34a-5p and hsa-mir-155-5p can be a potential diagnostic and therapeutic target for RA through various therapeutic strategies such as miRNA-Based Gene Therapy and Exosome-Derived miRNA (Chang et al., 2022; Peng et al., 2023; Zhang et al., 2022).

Additionally, an immune response is a reaction that happens within an immune system of an organism to fight against external factors such as pathogens, toxins, and viruses (Tu et al., 2021b). However, in RA patients, the immune system mistakenly activates autoimmune responses that attack the body’s own cells and cause chronic joint inflammation (Jang et al., 2022). As shown in Table 4.6.1, the BP of RPS27A that can trigger an immune response in RA includes an innate immune response, cytokine-mediated signalling pathway, and I-kappaB kinase / NF-kappaB signalling. The dysregulation of RPS27A can significantly contribute to the progression of immune response in RA. For instance, the RPS27A regulates gene expression of other genes involved in immune responses such as IL6, and CX3CL1 (Khayer et al., 2020). Hence, overexpression of RPS27A can cause dysregulation of gene expression of IL6, and CX3CL1 triggering aggressive immune responses such as chronic joint inflammation and cartilage degradation (Khayer et al., 2020). Moreover, RPS27A is responsible for the structural constituent of ribosomes, protein binding, and metal ion binding for MF. As RPS27A is a ribosomal protein, it regulates ribosome functions, especially protein synthesis and translation (Lari et al., 2021). Therefore, dysregulation of RPS27A can cause alteration in ribosome functions, aberrant synthesis of protein, and high production of inflammatory cytokines which can eventually lead to damage in joint tissues in RA (Lari et al., 2021).

Besides that, RPS27A are located in intracellular, cytosol, and ribosomes for CC. Overexpression of RPS27A can cause survival of immune cells, synoviocytes, and cartilage apoptosis developing RA (Khayer et al., 2020). Similarly, RPS27A is also enriched in the ribosome pathway to maintain biogenesis and function of the ribosome. Abnormal activation of this gene can lead to alteration in the biogenesis and function of ribosomes, contributing to the development of RA (Khayer et al., 2020).

On the other hand, based on Table 4.6.2, TP53 is enriched in T-cell proliferation involved in immune response, B-cell lineage commitment and innate immune response for BP. For instance, for T-cell proliferation involved in immune response, TP53 encodes p53 protein which regulates the T-cell proliferation and apoptosis (Ebrahimian et al., 2023). However, downregulation of TP53 is observed in RA that can cause impaired apoptosis and excessive T-cell proliferation triggering autoreactive immune response and chronic inflammation (Ebrahimian et al., 2023). Moreover, p53 binding, ubiquitin protein ligase binding, and damaged DNA binding are some of the MF enriched by TP53. For example, for the p53 binding, p53 binds with and triggers many important signalling pathways which is essential to suppress the adverse immune response in RA. However, low expression of TP53 can alter the p53 binding by allowing it to bind with TANK-binding kinase 1 (TBK1) which is implicated in the recognition of foreign pathogens (Shi & Jiang, 2021; Zeng et al., 2023). This ultimately causes the immune cells to not be able to distinguish between self-antigens and foreign pathogens causing it to attack self-antigens, self-tissues in RA.

Furthermore, chromatin, nuclear chromatin and nucleus are the CC of TP53 while Wnt signalling pathway, apoptosis pathway, MAPK signalling pathway are the KEGG pathway of TP53. For instance, the TP53 negatively activates the Wnt signalling pathway but in RA patients downregulation of TP53 can cause aberrant activation of Wnt signalling pathway, contributing to activation of immune cells such as T-cells, macrophages, and production of pro-inflammatory cytokines (Cici et al., 2019; Ding et al., 2023).

Therefore, current drug discovery needs to focus on modulating the immune response of RA to suppress inflammation and prevent long-term disability. Drugs with minimum to no side effects are essential for RA as the current medications are exhibiting adverse effects (Gupta et al., 2022; Radu & Bungau, 2021). Thus, *C.paniculatus* is a plant that is multifaceted and enriched in therapeutic properties against RA. Research supports the tendency of phytochemical constituents of *C.paniculatus* such as alkaloids, sterols, triterpenoids, polyalcohol, and sesquiterpenes contribute to the anti-depressant, and anti-inflammatory properties, suggesting that it can be a potential therapeutic target for RA (Fatima et al., 2021; Kothavade et al., 2015; Sharma et al., 2021; Yu et al., 2012).

*C.paniculatus* exhibits strong antioxidant properties proved by its ability to scavenge free radicals, inhibit peroxynitrite, and decrease the generation of oxygen species (Arya et al., 2022). The antidepressant potential is also exhibited by *C.paniculatus* as it shows alleviation in anxiety and depression behaviour of depression-induced mice (Chahuan et al., 2022). Therefore, *C.paniculatus* has the ability to treat the key features of RA such as inflammation, oxidant, and depression proving it can also be a potential therapeutic target for RA Fatima et al., 2021; Kothavade et al., 2015; Sharma et al., 2021; Yu et al., 2012). Hence, autodocking was performed between targeted proteins that are closely associated with RA and *C.paniculatus* phytochemical compounds to evaluate their tendency to act as drug candidates against RA.

Based on Figure 4.4.1, the upregulated and downregulated targeted proteins that are highly associated with RA are RPS27A and TP53 respectively as they play key roles in the progression of RA. Studies have revealed that abnormal expression of RPS27A can cause high production of the Ubiquitin protein leading to chronic joint damage due to its dysregulation (Behl et al., 2020). On the other hand, TP53 significantly regulates the matrix metalloproteinase expression in fibroblast-like synoviocytes (FLSs), and the proliferation of FLSs (Taghadosi et al., 2021). Dysfunction of TP53 leads to an elevated rate of inflammation and hyperproliferation of FLSs (Taghadosi et al., 2021). Consequently, it contributes to joint destruction, synovial hyperplasia, and treatment resistance for RA (Taghadosi et al., 2021). Thus, discovering drugs targeting RPS27A and TP53 is crucial to managing RA.

Hence, RPS27A and TP53 targeted proteins were docked with 20 compounds of *C.paniculatus* using Autodock Vina v1.5.7. The docking results showed that oleic acid exhibited the strongest binding affinity with RPS27A as is proven by its binding affinity compared to other ligands.(Table 4.8.3). This indicates a strong binding interaction between oleic acid and RPS27A as binding affinity is inversely proportional to the activity of the compounds (Al-Khayri et al., 2023; Enmozhi et al., 2020; Rizvi SM et al., 2013). On the other hand, zeylasterone showed the most effective binding interactions with TP53 among the other ligands. This shows that it has a higher activity and a potential to interact with TP53, suggesting that it can activate TP53 to encode p53 protein to suppress inflammation pathways.

The drug-likeness of oleic acid and zeylasterone was evaluated based on the five Lipinski rule which was proposed by Christopher A Lipinski in 1997 to examine the compounds’ physicochemical properties (Sharma et al. 2018). The Lipinski rule of five includes having a molecular weight of less than 500kDa so that it will not be a bulky molecule. This will not affect the permeability of the compounds as well (Al-Khayri et al., 202b; Variya et al., 2016). Moreover, a compound should have less than 5 lipophilicity which is a good indicator of its potential to penetrate biological membranes (Al-Khayri et al., 2023; Sharma et al. 2018). In addition, based on the Lipinski rule, the number of hydrogen bond donors and acceptors should be less than 5 and less than 10 respectively (Chang et al., 2004; Ertl et al., 2000). Violation of less than or equal to two parameters of Lipinski rule is acceptable (Ibrahim et al., 2020). The drug-likeness result revealed that both oleic acid and zeylasterone comply with Lipinski’s rules. (Table 4.9.1)

The major ADMET properties such as CaCo-2 permeability, Pgp-inhibitor, Pgp-substrate, Human Intestinal Absorption (HIA), Plasma Protein Binding (PPB), Blood-brain barrier (BBB), CYP inhibition, AMES toxicity, rat oral acute toxicity, and carcinogenicity were analysed. As presented in Table 4.9.2, the results stated that only oleic acid is CaCo-2 permeable whereas zeylasterone is not CaCo-2 permeable. The human colon adenocarcinoma cell lines (CaCo-2) act as human intestinal cell lines due to their similar morphology and function (Sharma et al. 2018). Thus, CaCo-2 acts as a substitution approach for human intestinal epithelium. Moreover, the inhibitor of P-glycoprotein (Pgp) activity leads to accumulation of drugs within the cells and causes cytotoxicity of the cell whereas Pgp-substrate serves as a barrier of drug absorption in the intestine (Callaghan et al., 2014). Both the compounds are not Pgp-inhibitor and Pgp-substrate, proving their characteristics as a good therapeutic drug.

The Human Intestinal Absorption (HIA) of both compounds is reported to be high which allows fast absorption of the drug as HIA is a crucial index to evaluate the oral bioavailability and intestinal absorption of the drug. In addition, the Plasma Protein Binding (PPB) is one of the most crucial mechanisms in drug distribution as it has the potential to influence oral bioavailability directly(Sharma et al. 2018). However, both oleic acid and zeylasterone have a high protein bound with a value of above 90% indicating the necessity for further development such as structure modification before it is used as a therapeutic drug against RA.

Moreover, identifying the Cytochrome P450 (CYP) inhibition is also crucial as it is a major route for elimination of many drugs and prevents drug-drug interactions (DDIs) (Zhao et al., 2021). Based on Table 4.9.3, the major CYP isoforms such as CYP1A2, CYP2C19, CYP2C9, CYP2D6, and CYP3A4 are identified for oleic acid and zeylasterone. The CYP1A2 is involved in metabolism of theophylline while CYP2C19 is involved in proton pump inhibitors and antiplatelet drugs metabolism(Zhao et al., 2021). CYP2C9 is implicated in the metabolism of NSAIDs (Zhao et al., 2021) whereas CYP2D6 is responsible for drug metabolism such as antidepressant drugs and oxidative reactions (Taylor et al., 2020). Additionally, CYP3A4 is involved in metabolism of immunosuppressants and statins (Zhao et al., 2021). It is found that both the oleic acid and zeylasterone are non-inhibitors for all CYP isoforms. This proves that they will not lead to impaired biotransformation of drug and drug-drug interactions. Hence, they will not cause any issues with CYP-mediated metabolism during drug development.

Another important aspect of the PreADMET is the toxicity of a drug. The Ames Mutagenicity Assay (AMES) and carcinogenicity test were performed. They evaluate the mutagenicity, and carcinogenicity of the compounds to avoid being toxic. Based on PreADMET results, both compounds are non-mutagens and non-carcinogens. In addition, the rat oral acute toxicity was also analysed for safety evaluation. The result indicated that oleic acid and zeylasterone are safe to be drug candidates. The human-ether-a-go-go-related gene (hERG) blockers are crucial to determine and eliminate the compounds that have the potential to cause fatal cardiac arrhythmias (Mackenzie, 2018). The result showed that both compounds are hERG blockers. Hence, oleic acid and zeylasterone are potential drug candidates but also require alteration of compound structures to optimise their stability and safety to use as drugs.

## CONCLUSION

In summary, this research indicates that analysing the top hub genes, immune response, and miRNA-hub genes networks associated with RA via integrated bioinformatics analyses helps to comprehend the mechanism of RA progression in-depth and provide alternative therapeutic approaches against RA. Besides, this study identified RPS27A, UBB, UBC, UBA52, PSMD4, PSMD1, PSMD7, PSMB7, PSMD8, PSMA7 as top 10 upregulated hub genes associated with RA whereas TP53, ACTB, AKT1, GAPDH, CTNNB1, EGFR, TNF, IL6, MYC, ANXA5 as the 10 top-most downregulated hub genes. Moreover, analysing the miRNAs associated with RA and their biological role in the pathogenesis of RA provides insights into discovering miRNA-based diagnostics and therapies. The miRNA-hub genes network identified hsa-mir-23b-3p as highly associated with the top 10 upregulated hub genes whereas hsa-mir-34a-5p and hsa-mir-155-5p with the top 10 downregulated hub genes. They have the potential to act as both biomarkers and therapeutic targets against RA. In addition, the oleic acid and zeylasterone compounds from *C.paniculatus* have a high tendency to serve as drug candidates as they exhibit a strong binding affinity with targeted proteins and pass the druglikeness and PreADMET tests. Hence, the immune responses of RPS27A and TP53 were revealed and it is proven that miRNA, phytochemical compounds of *C.paniculatus* have therapeutic effects against Rheumatoid Arthritis.

## ACKNOWLEDGEMENTS

The researchers would like to thank Management and Science University for supporting the completion of this research.

